# Structural basis of Arf1-driven membrane tubulation

**DOI:** 10.64898/2026.01.22.700998

**Authors:** Caroline Haupt, Dmitry A. Semchonok, Ambroise Desfosses, Sebastian Daum, Sarah Neudorf, Farzad Hamdi, Panagiotis L. Kastritis, Milton T. Stubbs, Kirsten Bacia

## Abstract

Membrane tubules form at Golgi compartments to facilitate membrane and cargo flow in intracellular trafficking. Here we show that the small GTPase Arf1, an inducer of membrane curvature and key regulator of trafficking, is able to form strongly curved tubules in the presence of lipids and GTPγS *in vitro* without the need for further coat components. Using cryo-electron microscopy, we determined the structures of tubular Arf1-scaffolds with diameters of 195 and 215 Å at 3.1 and 3.8 Å resolutions, respectively. The nucleotide-bound globular domains of Arf1 form polar helical lattices (*i.e.* directional assemblies with distinct start/finish orientations), with conserved interfaces and a consistent back-to-face orientation along the filaments. The rigid coat is tethered to the membrane by a flexible linker and anchored by an amphipathic helix (AH) that is free to diffuse and make space within the leaflet, allowing for accommodation of transmembrane cargo. The diversity of tubular diameters observed would allow various cargo sizes to be accommodated in the lumen, while maintaining the local coat architecture. Apart from serving as tubular transport intermediates, Arf1-scaffolds may also play a role at the neck of COPI vesicle on the route to scission.

## Introduction

Intracellular transport is essential for maintaining homeostasis in eukaryotic cells. Proteins are transported between organelles primarily by two forms of membrane carriers: vesicles and tubules^1^, both of which require the generation of high membrane curvatures, i.e. small diameters. Such small diameters are achieved through polyphilic protein-membrane and protein-protein interactions. Prime examples of such processes include the COPI and COPII vesicular transport systems, which generate vesicles of 200 - 500 Å^2^ or 600 - 1000 Å^3,4^ in diameter, respectively, via the combined action of (i) amphipathic helix (AH) leaflet insertion and (ii) anchoring of a curved protein scaffold^5–7^.

In eukaryotic cells, the Golgi complex functions as a pivotal intracellular sorting hub. Arf1, a member of the Sar1/Arf1 family and the Ras superfamily of GTPases^8^, is essential for maintaining Golgi structure and regulates many trafficking processes associated with Golgi compartments. Arf1 is highly conserved and interacts with a large number of regulators and effectors, recruiting coat protein complexes. Like other Ras superfamily members, Arf1 transitions between a GDP- and a GTP-bound state^9^. Arf1 is N-terminally myristoylated (myr-Arf1), and in the GDP-bound state the myristoyl moiety is sequestered within a hydrophobic cavity of the protein (Fig. 1a)^10^. In the GTP-bound state, major structural rearrangements result in restructuring of the N-terminal peptide to form an AH and exposure of the myristoyl group that allows insertion into the phospholipid leaflet (Fig. 1b)^11^, anchoring the GTP-bound globular (G-) domain to the membrane.

**Fig. 1:**
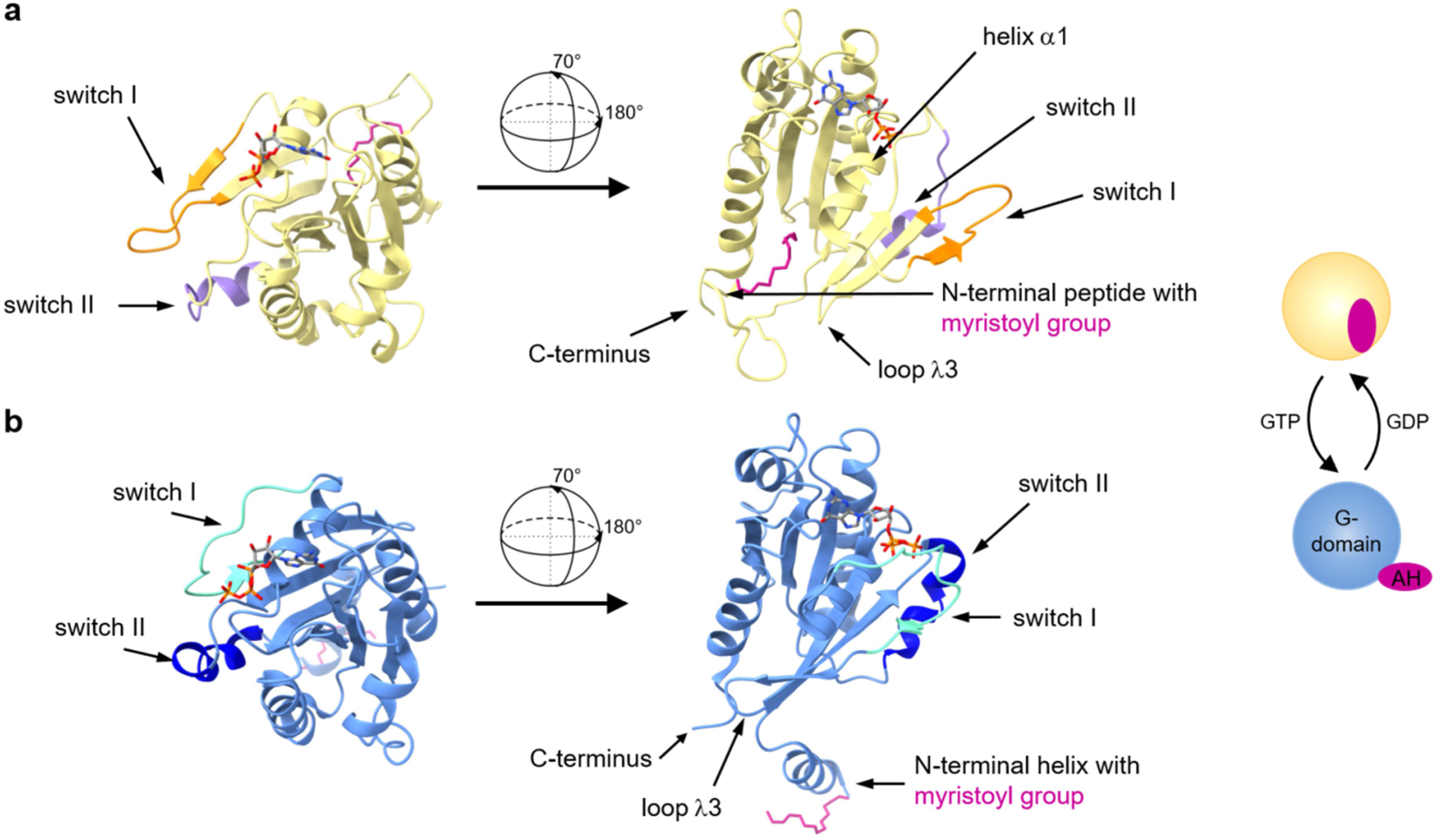
Structures of myr-Arf1 in solution. **a** GDP-bound myr-Arf1 (yellow, PDB: 2k5u)^10^. Structural features of the small GTPase GDP-GTP switch are highlighted (switch I: orange, switch II: purple, interswitch loop λ3). Arf1 consists of a central β-sheet surrounded by several α-helices that together form the globular G-domain. The secondary structure nomenclature follows that for Ras proteins^86^. The N-terminal peptide is unstructured, whereas the covalently linked myristoyl group (pink) is buried in a hydrophobic cleft in the GDP-bound state. GDP and the myristoyl group are shown in stick representation. **b** GTP- and membrane- (bicelle-) bound myr-Arf1 (blue, PDB: 2ksq)^11^. The same structural features are highlighted (switch I: cyan, switch II: dark blue), with GTP and the myristoyl group (pink) shown in stick representation. The switch regions undergo significant conformational changes upon exchange of GDP by GTP, deforming the hydrophobic cleft that harbors the myristoyl moiety in the GDP-bound state. This exposes the myrAH in the GTP-bound state to allow anchoring the G-domain to the membrane^40^. Exposure and retraction of the N-terminal helix upon nucleotide exchange are also illustrated schematically, G-domain and AH are labeled.

Arf1 thus functions as a bimodal switch, whereby GDP/GTP state cycling is coupled to membrane recruitment, since (i) intrinsic Arf1 GTPase activity is low^12^ (in general requiring the presence of Arf GTPase activating proteins or GAPs), and (ii) Arf1 product release is slow (requiring the action of Arf guanine nucleotide exchange factors (GEFs) to exchange GDP for GTP)^13^. GEFs Gea1/Gea2 (yeast) and GBF1 (mammals) function at early Golgi compartments to mediate retrograde trafficking to the ER^14–16^, whereas Sec7 (yeast)^17^ and BIG1/BIG2 (mammals)^18^ are localized to late Golgi compartments, the *trans*-Golgi network (TGN) and recycling endosomes. ARFGAP1-3 and their yeast orthologues act at the early Golgi, whereas most other Arf GAPs function at the TGN, endosomes and plasma membrane^14^.

Membrane-anchored Arf1 has been shown to recruit coat complexes *in vivo* and *in vitro*^19,20^, and the structure of the COPI coat, stabilized by non-hydrolyzable GTPγS on liposomes *in vitro*^21^ as well as with GTP in cells^22^, has been elucidated through a combination of X-ray crystallography and cryo-electron tomography (cryo-ET). In addition to recruiting adaptor and coat proteins, Arf1 plays a key role in membrane fission by generating a highly curved tubular structure at the neck of COPI buds^23,24^, analogous to Sar1 at the neck of COPII buds^25^, and mutations in the AH of Sar1 and Arf1 affect both tubulation activity and vesicle scission^7,26^. Nevertheless, the mechanism underlying how Arf1 (or Sar1) interacts with the bilayer and with itself to drive membrane fission remains unclear, underscoring the necessity for high-resolution structural data.

*In vivo*, Arf1-positive tubes play a role in both antero- and retrograde trafficking at the Golgi, potentially functioning as transport intermediates^27^. Bottanelli *et al.*^27–29^ have proposed that these intracellular tubules result from an Arf1-induced bilayer tubulation process akin to that observed in *in vitro* reconstitution studies and highlighted that Arf1-positive tubules constitute a significant component of Golgi-derived membrane flow. From the *in vitro* reconstitution experiments it was concluded that the generation of a high membrane curvature was linked to Arf1 dimerization^5,30^.

Motivated by the hypothesized physiological relevance of Arf1 tubular assemblies at COPI bud necks and their potential role as tubular transport intermediates *in vivo*, we have applied cryo-EM helical reconstruction to analyze the morphology and structure of *in vitro* reconstituted Arf1 assemblies. Our findings provide structural insight into the adaptability of Arf1 scaffolds across diverse curvatures, shedding light on the molecular basis of Arf1-mediated membrane remodeling.

## Results

### Structure of the myr-Arf1-coated membrane tubules

Incubation of giant unilamellar vesicles (GUVs) with myr-Arf1 (hereafter referred to as Arf1) and the non-hydrolyzable GTP analogue GTPγS resulted in the formation of long, rigid and narrow Arf1-covered membrane tubules (Supplementary Fig. 1a-c, Supplementary Movie 1a, b). The tubulation reaction was plunge-frozen on cryo-EM grids and assessed by cryo-fluorescence microscopy (Supplementary Fig. 1d, e). Visual inspection of the cryo-EM micrographs revealed tubules with a broad distribution of diameters ranging from 150 to 400 Å (Fig. 2a, Supplementary Fig. 2). Two-dimensional (2D) classification of tubule projections revealed well-populated class averages within the diameter range of 180 to 280 Å (Fig. 2b, schematic representation of the data analysis workflow in Supplementary Fig. 3). Fourier analyses of the 195 and 215 Å diameter 2D class averages (power spectra in Supplementary Fig. 4) demonstrate a regular arrangement within the membrane tubules, indicating helical arrays of Arf1 assembled on the lipid bilayers. The helical symmetries of the different diameter populations were determined by analysis of the respective average power spectra (Supplementary Figs. 4, 5), and validated by the appearance of high-resolution features in the resulting 3D reconstruction after helical refinement.

**Fig. 2:**
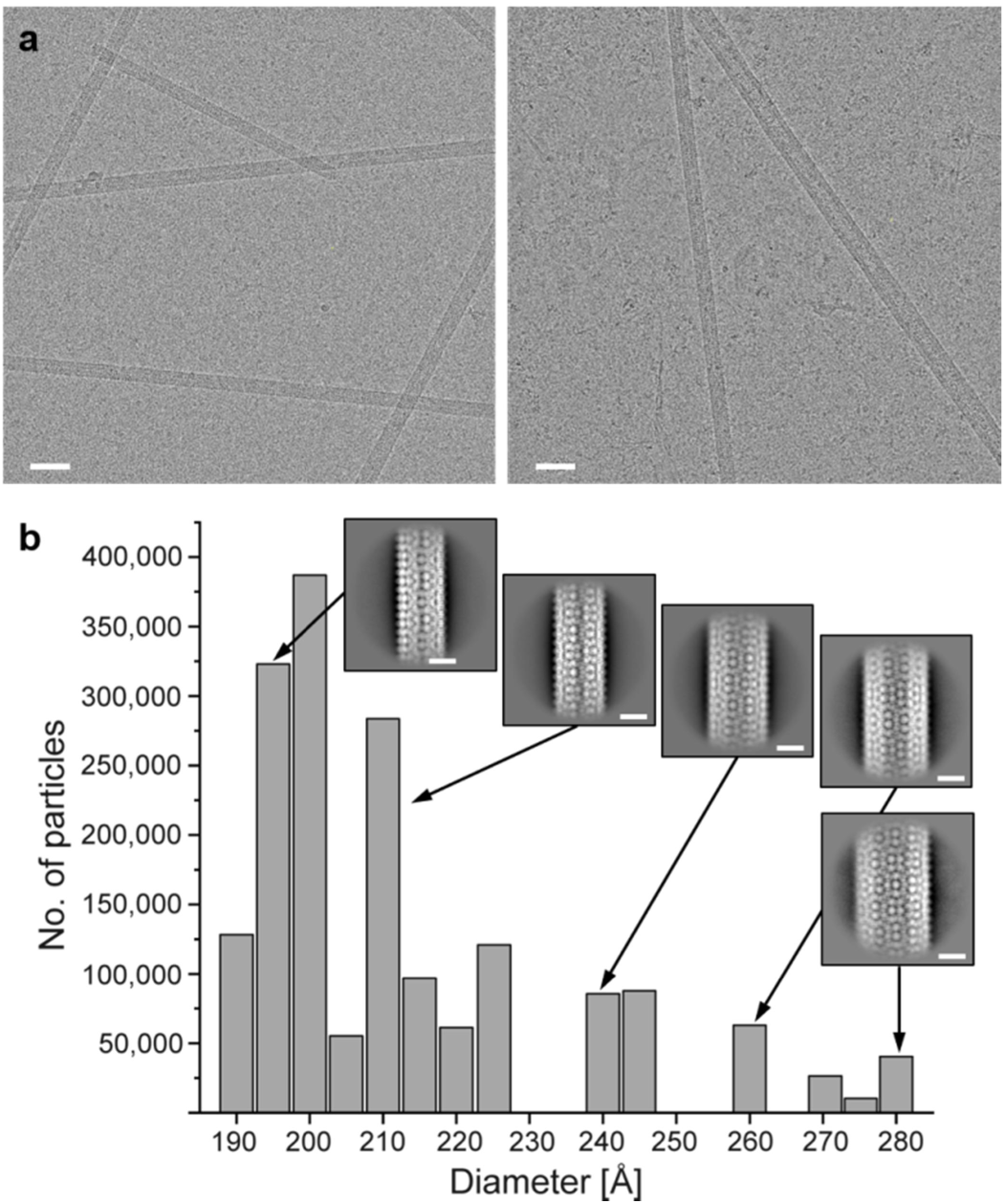
Arf1-mediated membrane reshaping. **a** Representative cryo-electron micrographs of Arf1-decorated tubules highlighting the large variability in tubule diameters. Scale bars: 500 Å **b** Size distribution of tubule diameters with representative 2D class averages. The most abundant class of 200 Å diameter tubules was excluded from further analysis due to large heterogeneities in the corresponding power spectra. The displayed numbers of particles result from the particle picking/extraction procedure described in the Methods section (12,483 raw movies). Scale bars: 100 Å

Among the observed tubules, two dominant populations with diameters of 195 Å and 215 Å were analyzed in detail, representing 27% and 7% of the final 2D classification set, respectively (Supplementary Fig. 3). Helical reconstruction of the 195 Å tubules revealed a right-handed two-start helix with about 2 × 16 monomers per turn (Fig. 3b), whereas the 215 Å tubules form a right-handed three-start helix with about 3 × 18 monomers per turn (Fig. 3f). The tubules are polar, resulting in an A- and a B-end (Fig. 3b, f). High-resolution 3D reconstructions up to spatial resolutions of 3.1 Å (195 Å diameter tubules) and 3.8 Å (215 Å diameter tubules), as determined by the 0.143 Fourier shell correlation (FSC) criterion^31^ (Supplementary Fig. 6a, b), could be achieved, with helical parameters summarized in Supplementary Table 1.

**Fig. 3:**
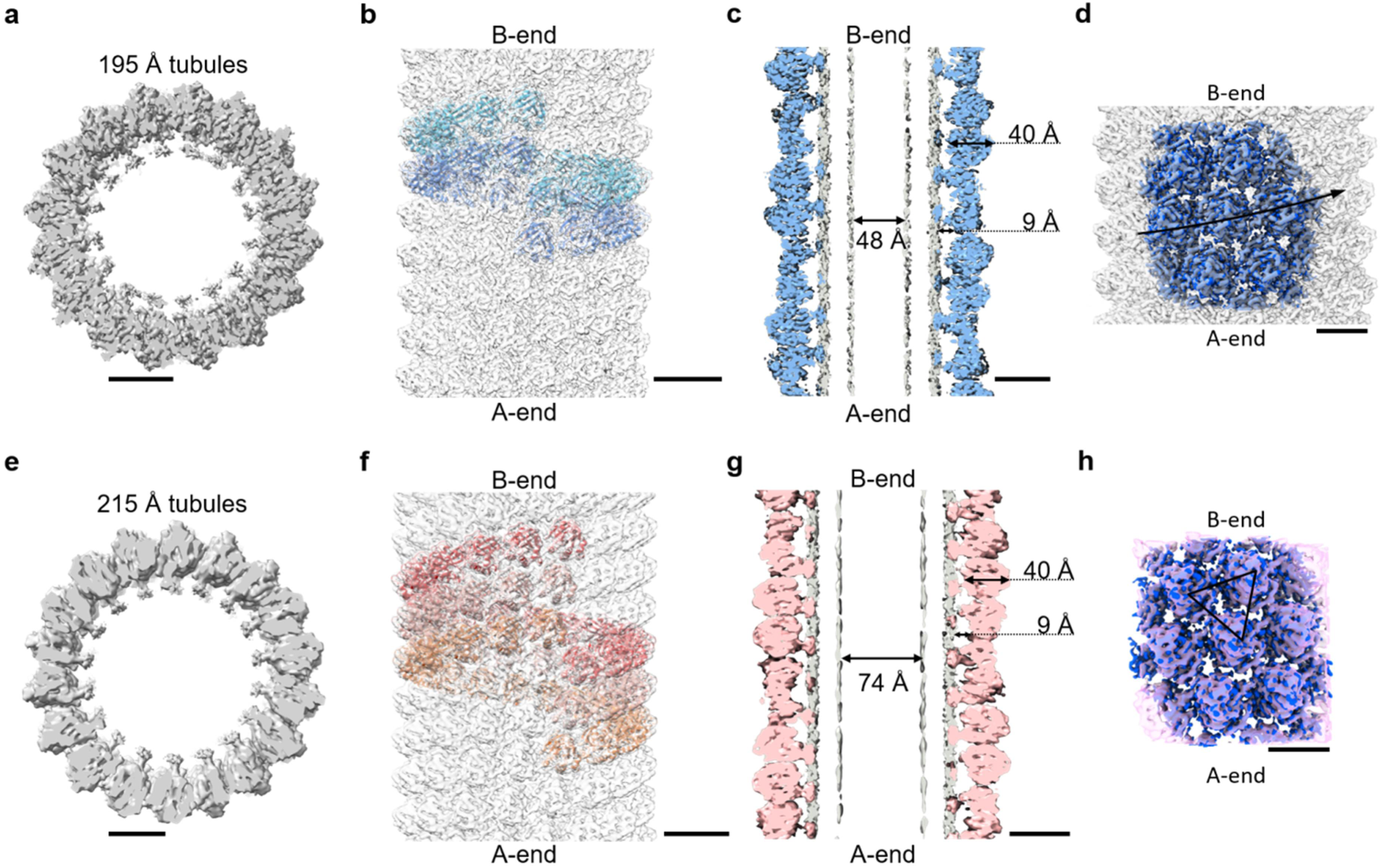
**Cryo-EM structure of the Arf1-coated membrane tubules.** Cryo-EM 3D reconstructions of the Arf1 helical assembly of the 195 Å diameter tubules resolved at 3.1 Å resolution (**a**-**d**) and of the 215 Å diameter tubules resolved at 3.8 Å resolution (**e**-**g**). **a**, **e** Axial views of central slices along the tubules. **b**, **f** Side views of the tubules with cryo-EM maps overlaid as transparent volumes. Start and finish ends of the polar (directional) tubules are labeled with A- and B-ends, respectively. Asymmetric units related by helical symmetry are shown in a single color as cartoon representations, whereas alternative colors represent asymmetric units related by rotational symmetry Cn (C2 for 195 Å tubules (**b**), C3 for 215 Å tubules (**f**)). **c**, **g** Central slice side views of cryo-EM density maps showing the protein layer, membrane inner lumen, and the protein-membrane distance. A composite map, sharpened and filtered independently to distinguish membrane (grey) and protein (blue for 195 Å (**c**); light pink for 215 Å (**g**)), is shown. **d** Zoomed-in view of the reconstructed cryo-EM 3D density map of the Arf1 helical assembly of the 195 Å diameter tubules (in grey) is shown with the extracted density for modelling an Arf1 molecule surrounded by six neighboring Arf1 molecules in blue. The arrow indicates the progression of a single helical filament. **h** Zoomed-in view of the overlay of the 195 Å (blue) and 215 Å (pink) tubule cryo-EM maps demonstrating conserved lattice structures. Black solid lines indicate the triangular arrangement for the three interfaces. Scale bars: 50 Å

The local resolution of the final 3D maps varies with the radius of the tubules, with the G-domain exhibiting the highest resolution (Supplementary Fig. 6c, d). Regions of high contrast inside the tubes (Fig. 3c, g) delineate the phosphate headgroup positions of the lipid bilayer, with a nominal thickness of 27 Å; the slightly higher density of the outer leaflet is consistent with partial embedding of the N-terminal amphipathic helices as seen in molecular dynamics (MD) simulations of monomeric membrane-bound Arf1-GTP^32^. In the tubes, the distance from the AH C-terminus to K16 (the first ordered residue of the G-domain) of approximately 9 Å is considerably longer than in the MD simulations, where the G-domain appears to contact the membrane (Supplementary Fig. 7a). The formation of the tubular Arf1 coat, with radially distributed forces, results in minimizing the degrees of freedom of single Arf1 G-domains to keep a distinct distance of the Arf1 scaffold to the membrane while maintaining two-dimensional degrees of freedom of the AH in the membrane.

A comparison of the two tubule reconstructions shows that although their inner lumen diameters differ, the thicknesses of their outer protein layers are similar (Fig. 3c, g). Both reconstructions also exhibit the same lattice arrangement of the Arf1 G-domains (Fig. 3h), as seen in the unrolled lattices of the cylindrical 3D EM maps (Supplementary Fig. 8a, b). Similarities in the averaged power spectra of 2D class segments from tubules of other diameters suggest that the overall lattice structure is conserved in each of our tubule preparations (Supplementary Figs. 4, 8). Processing of a third 240 Å diameter tubule also revealed the same lattice arrangement within a four-start helical structure with about 4 × 21 monomers per turn (Supplementary Fig. 9), with the same lattice arrangement, although we were unable to refine the structure to high resolution, possibly due to inhomogeneities within the class averages.

The quality of the 195 Å diameter tubule reconstruction allowed the complete G-domain (K16 to K178) to be built, together with the GTPγS nucleotide analogue and a Mg^2+^ ion coordinated by the sidechains of T48 and T31 and the β- and γ-phosphates of the nucleotide (Fig. 4a). The final Arf1 model exhibited a root mean square deviation value of 0.639 Å with the crystal structure of Arf1^33^ (Supplementary Fig. 10a), and could readily be fitted to the lower-resolution cryo-EM density map of the 215 Å tubules with minimal adjustments (Fig. 3f).

**Fig. 4:**
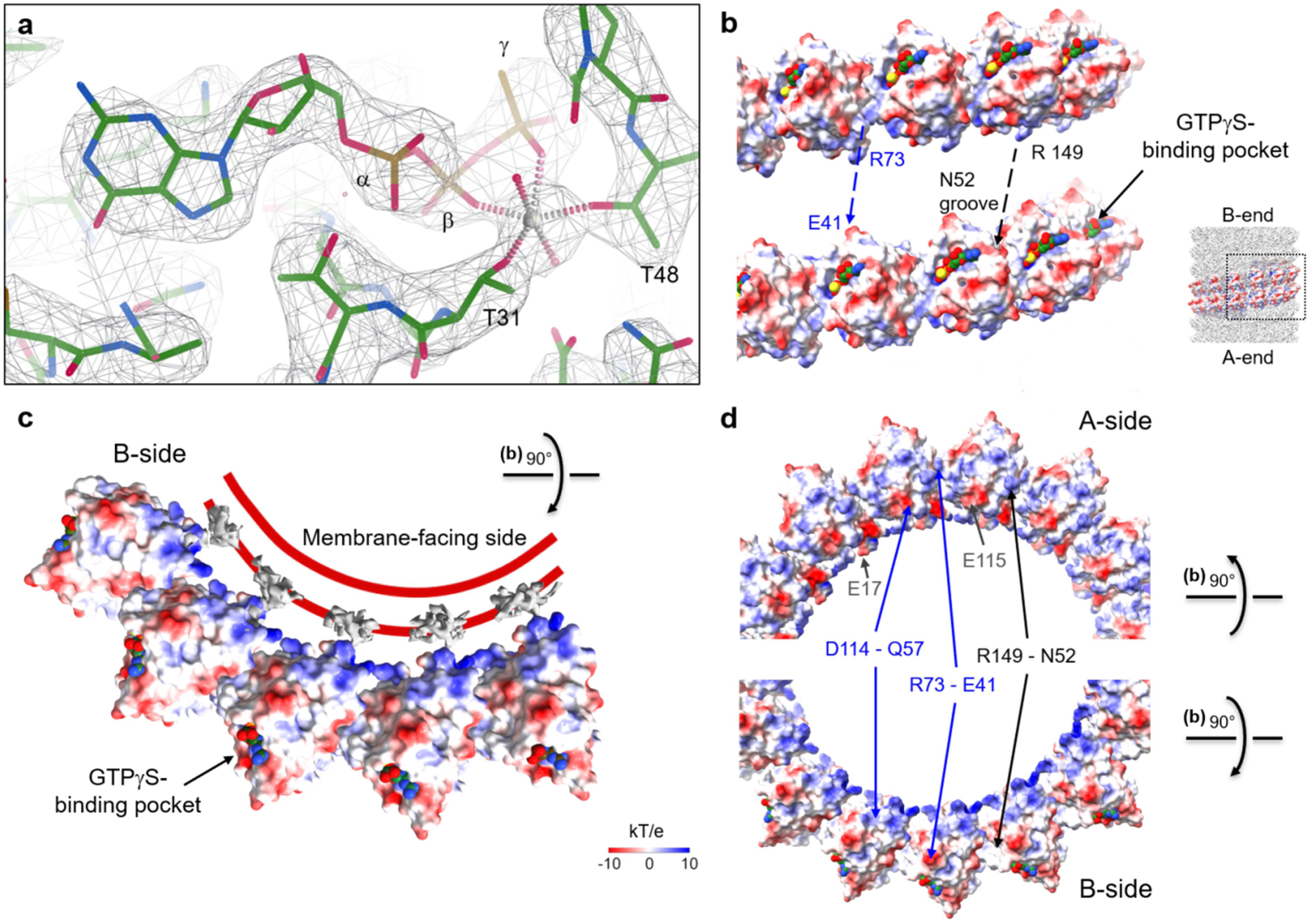
Structure of Arf1 in the tubular membrane assembly. **a** The high resolution of the cryo-EM density map (mesh representation) allows fitting of the nucleotide GTPγS and magnesium ion to the active site. The map is in agreement with an octahedrally coordinated magnesium ion as was reported in the crystal structure of mouse Arf1 (PDB: 1o3y)^33^. **b-d** Electrostatic surface potential maps (in kT/e) of individual filaments highlight the role of charge-charge interactions in stabilizing the Arf1 helical assembly (**b**, **d**), with the basic interior surface of the protein coat facing the negatively charged headgroups of the membrane outer leaflet (**c**). Side view (**b**) and axial view (**d**) of two adjacent filaments, separated to better illustrate the interaction sides, reveals that the basic residues R149_2a_ and R73_3a,1ʾ_ are prominently exposed at inter-filament interfaces 2 and 3 (black and blue labels, respectively), juxtaposed by acidic residues (grey labels). The corresponding location of the two filaments within the tubule is indicated. Panel (**c**) shows five neighboring Arf1 molecules along the helical filament. The extracted density of the unmodeled N- terminal peptide is shown in grey. The curved membrane bilayer is indicated (red solid lines) with the GTPγS-binding pocket facing outward.

The nucleotide-binding site is located on the exterior of the tubules (Fig. 4b, c), rendering it accessible to GAP proteins. While the G-domain was well defined in the map, density for the N-terminal amphipathic helices was lacking, which may simply be a result of them not following the G-protein coat symmetry, or might reflect a more general fluidity within the membrane outer leaflet. Residual density protruding from the protein layer into the outer leaflet of the membrane, spanning a distance of approximately 9 Å (Fig. 3c, g), can be attributed to an extended linker peptide connecting the G-domain and the AH, although it is not well defined. The orientation of this membrane-proximal density perpendicular to both the tubule axis and the proposed position of the AH (Fig. 3c, g), is consistent with solid-state NMR and MD simulation experiments^32^ (Supplementary Fig. 7a-c).

### G-domain protein-protein interactions within the Arf1 helical scaffold

In the 195 Å diameter tubule G-domain coat, each Arf1 subunit is surrounded by six neighbors with three distinct asymmetric interfaces (designated as 1, 2 and 3) and opposing interacting surfaces (named a and b) (see Fig. 5a). Interface 1 (half interface area 454 Å^2^, according to PISA server)^34^ occurs between consecutive Arf1 molecules along a helical row, whereas interfaces 2 (half area 232 Å^2^) and 3 (half area 355 Å^2^) are found between adjacent filament layers (Fig. 5a). Due to a similar lattice arrangement (Fig. 3h), corresponding interfaces are observed in the 215 Å tubules.

**Fig. 5:**
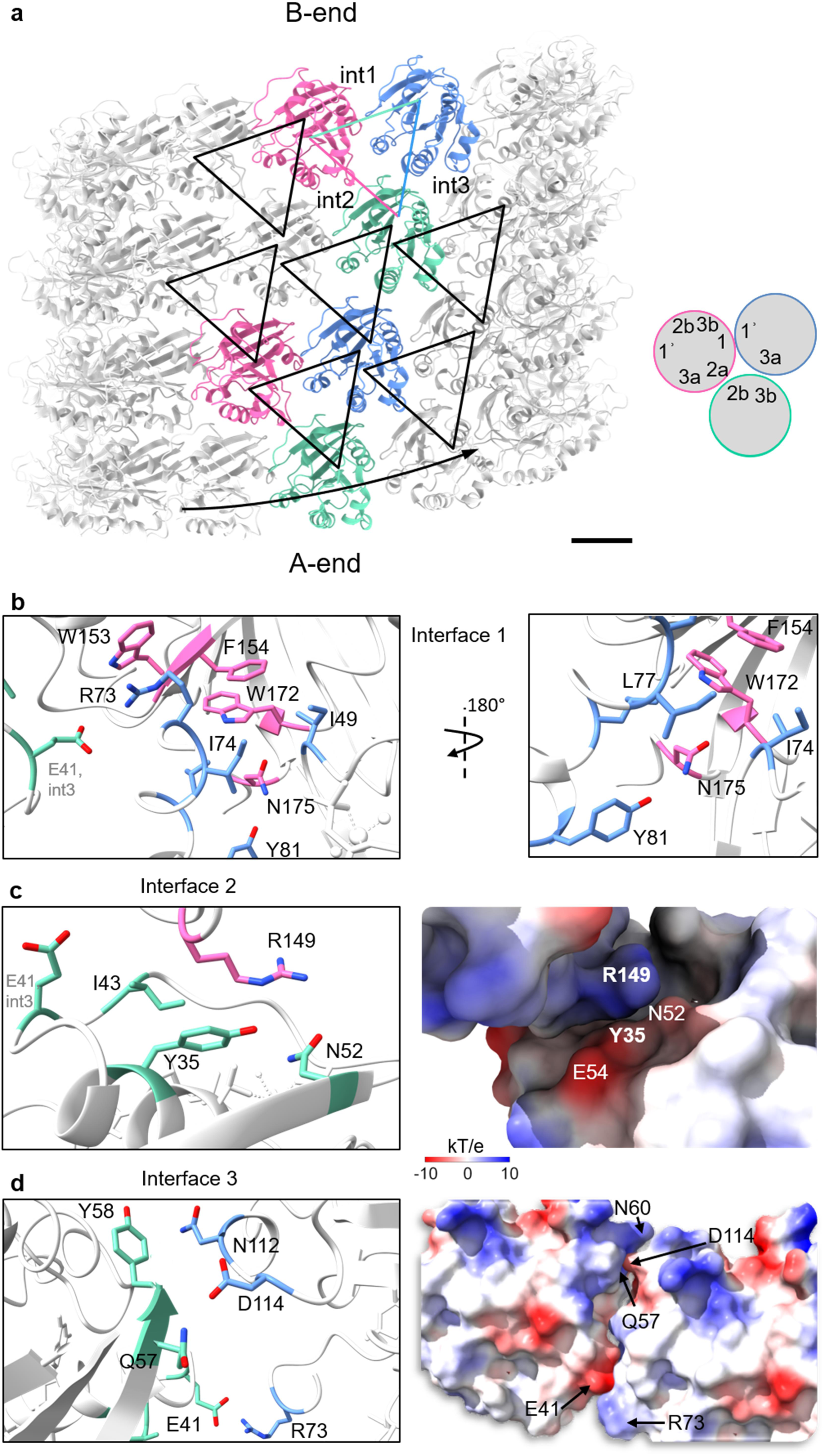
Close-up view of the three distinct interfaces. **a** A schematic diagram of the repetitive triangular interaction scheme within the Arf1 helical assembly with intermolecular contacts along the helical filament (interface 1, marked as int1) and contacts between adjacent filaments (interfaces 2 and 3, marked as int2 and int3, respectively). Two triangular molecular arrangements are highlighted. Repeated interfaces are marked with black lines, and filament progression is indicated by the arrow. A simplified scheme illustrates the nomenclature of the interfaces of single Arf1 molecules, where 1 and 1ʾ constitute the respective opposing sites for intra-filament interface 1, and a and b the respective opposing sites for inter-filament interfaces 2 and 3. Scale bar: 20 Å **b** Interface 1 forms between two consecutive Arf1 molecules along a helical row. This interface is primarily stabilized by hydrophobic interactions between F154_1_ and W172_1_ (1, pink) with I49_1ʾ_, I74_1ʾ_ and L77_1ʾ_ (1ʾ, blue). Additional cation-π-stacking (W153_1_ - R73_1ʾ/3a_) and hydrogen bonding (N175_1_ - Y81_1ʾ_) reinforce the interaction. Residues from 1ʾ (blue) belong to the switch II region. Non-interacting residues are shown in white. **c** Interface 2 forms between Arf1 molecules on neighboring filaments and is mediated by hydrogen bonds between residues of the switch I region (2b, green; I43_2b_, N52_2b_) and interactions between Y35_2b_ to R149_2a_. Left panel: Ribbon diagram with interacting residues shown in stick representation. Right panel: Electrostatic surface potential map of interface 2, highlighting the pronounced polar nature of the region. The loop containing I43_2b_ is positioned towards the back. Note the large cavity occupied by the arginine side chain R149_2a_ (2a, pink). **d** Interface 3 consists of inter-filament interactions. Residues from the interswitch region (loop λ3; 3b, green; Q57_3b_, Y58_3b_) form hydrogen bonds with D114_3a_ and N112_3a_ (3a, blue), while an electrostatic interaction between E41_3b_ and R73_3a,1ʾ_ further stabilizes the interface. Right panel: Electrostatic surface potential map of interface 3. The positions of key interacting residues are labeled. The D114_3a_ side chain is tightly packed, leaving no space for bulkier residues.

HADDOCK energy calculations^35–38^ yielded a negative score for each interface (Supplementary Table 2), indicating their thermodynamical feasibility. Although the absolute values are small for typical protein-protein interactions, the multiplicity of contacts afforded by the pseudo-crystalline G-domain arrangement, combined with AH membrane attachment as well as electrostatic attraction of the strongly positively charged inner surface (Fig. 4c) to the negatively charged headgroups of the membrane tubule outer leaflet, are likely to confer stability to the tubules. Protein coat interface 1 between consecutive Arf1 molecules along one helical strand (Fig. 5b) exhibits the largest number of intermolecular contacts, with equally balanced favorable van der Waals, electrostatic and desolvation energies (Supplementary Table 2). This interface is dominated by hydrophobic interactions between W172_1_ and F154_1_ on one Arf1 molecule (int 1) and I49_1ʾ_, I74_1ʾ_ and L77_1ʾ_ on the opposing interface (1ʾ, Fig. 5b), and further supported by hydrogen bonds (N175_1_ - Y81_1ʾ_) and cation-π stacking (W153_1_ - R73_1ʾ_). Cation-π stacking, between R149_2a_ and Y35_2b_, also contributes to the inter-filament interface 2, which due to its positive desolvation energy is rather weak, but is still notable considering the strong interaction per surface area (Fig. 5c). Basic side chains R73_1ʾ_,_3a_, R104, R109, and R149_2a_ contribute to a net positive charge on the A-surface of each filament that juxtaposes a negatively charged B-surface of the adjacent filament formed by E17, E41_3b_, and E115 (Fig. 4b, d).

Single-point mutations were made to probe the contribution of specific residues on membrane tubule formation (Fig. 6) and tested for membrane binding in a liposome flotation assay (Supplementary Fig. 11). The mutant Y35A_2b_ exhibits no tubulation activity and no membrane binding, consistent with previous studies showing a significant impact of tyrosine 35 on membrane tubulation^5,30^. Mutation of juxtaposing residue R149_2a_ to alanine, tyrosine or tryptophan reduces the *in vitro* tubulation activity (Fig. 6d-f) compared to the wildtype (Fig. 6a).

**Fig. 6:**
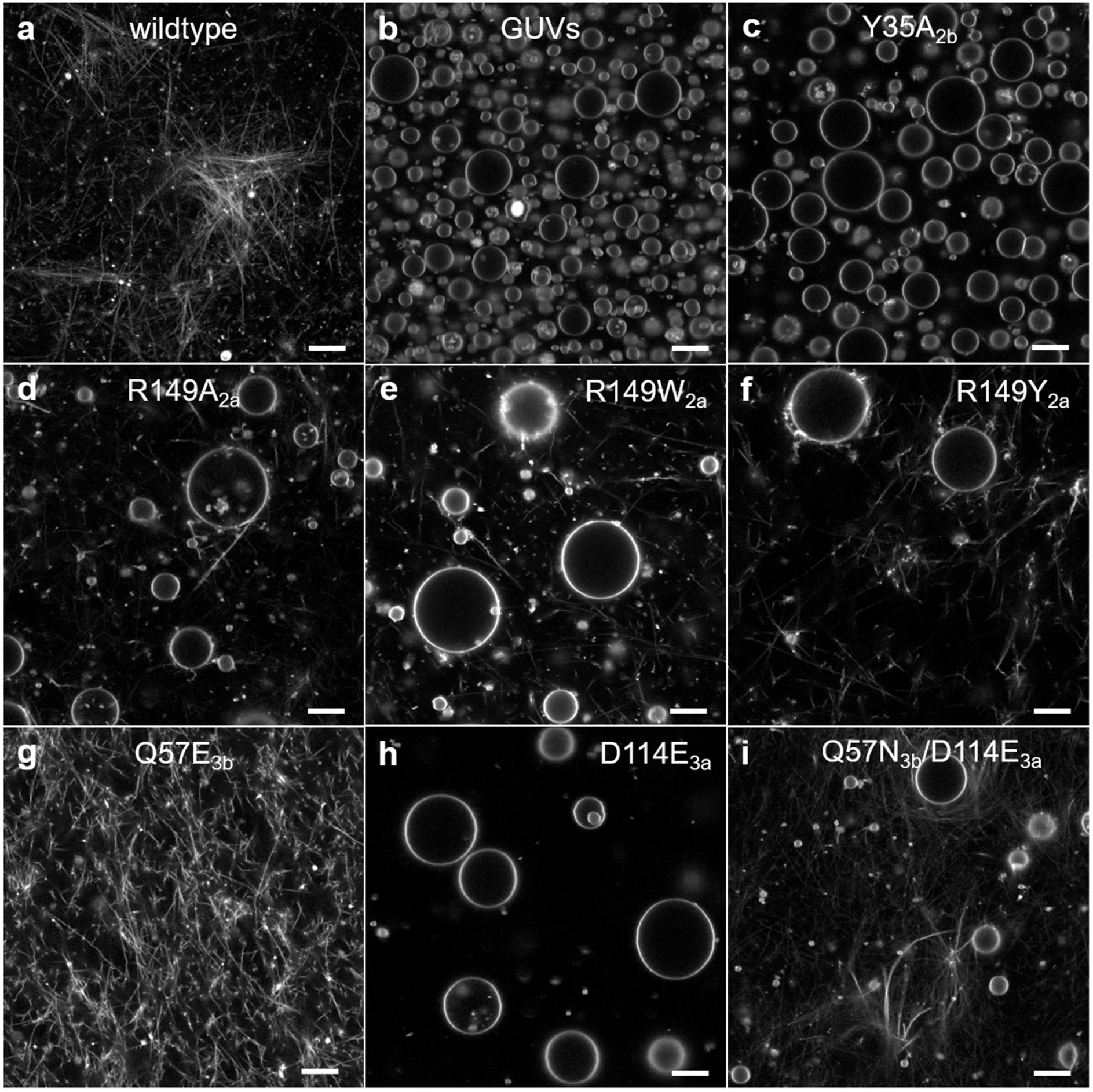
Membrane tubulation by Arf1 and Arf1 variants visualized by confocal microscopy. Fluorescently labeled membranes were incubated with wildtype Arf1 (**a**) and Arf1 variants R149A_2a_ (**d**), R149W_2a_ (**e**), R149Y_2a_ (**f**), Q57E_3b_ (**g**), and Q57N_3b_/D114E_3a_ (**i**). These conditions led to distinct membrane tubulation, forming a spacious network of straight, rigid tubules after 400 - 500 minutes. In contrast, incubation with the variants Y35A_2b_ (**c**) and D114E_3a_ (**h**) resulted in predominantly intact GUVs, indicating reduced tubulation. A control image of untreated GUVs is shown in panel (**b**). For a better visualization of the tubules, the images are shown at γ = 0.45 (nonlinear intensity scale). Scale bars: 10 μm

A salt bridge between E41_3b_ and R73_3a,1ʾ_ and hydrogen bonds between Q57_3b_ and D114_3a_ and between Y58_3b_ and N112_3a_ form the core of interface 3 (Fig. 5d). Interface 3 is heavily electrostatically driven, compensating for the unfavorable desolvation and weak van der Waals energies (Supplementary Table 2). Introducing an additional negative charge (Q57E_3b_) did not disrupt tubulation activity (Fig. 6g), but elongating the opposing chain (D114E_3a_) strongly inhibited tubulation (Fig. 6h) although membrane-binding was still detectable (Supplementary Fig. 11c). Tubulation activity was restored by replacing the opposing glutamine residue with the shorter asparagine residue (Q57N_3b_/D114E_3a_; Fig. 6i). Membrane flotation assays corroborate the tubulation activity results for all Arf1 variants (Supplementary Fig. 11).

Many residues involved in the interfaces are located in functionally relevant structural features of small GTPases (Fig. 1): I74_1ʾ_, L77_1ʾ_ and Y81_1ʾ_ belong to the switch II region and are involved in interactions with coatomer proteins^2,21,39^; I43_2b_ and N52_2b_ belong to the switch I region, constituting the interaction site for adaptor proteins that promote nucleotide exchange and enhance GTPase activity^16,40–43^; and residues Q57_3b_ and Y58_3b_ are part of the flexible interswitch loop λ3. Importantly, the interfaces are dependent on the G-domain adopting the GTP-bound conformation.

## Discussion

We have determined the structure of the activated small GTPase Arf1-GTPγS bound to membranes. The globular Arf1 G-domain encases a membrane tubule through self-polymerization into a helical scaffold, with the non-hydrolyzable GTP analog bound in a pocket on the outer surface of the tubules. Nucleotide binding induces the characteristic GTP-bound switch II conformation, which in the present structures is dedicated to intermolecular interface 1 interactions. A number of mutations in this region have been reported^30,44^ whose behavior in reconstitution experiments is consistent with a role in intra-filament interactions. These are complemented by our mutagenesis studies focusing on inter-filament interfaces. Single-point mutations of R149_2a_ have a moderate effect on tubulation, whereas mutants Y35A_2b_ (which juxtaposes R149_2a_ in interface 2) and D114E_3a_ impair tubulation. The G-domain is anchored to the membrane by an N-terminally myristoylated amphipathic helix (AH) via a flexible linker peptide^32^, giving the AH limited freedom to diffuse in the membrane outer leaflet. It has been demonstrated that lowering the amphiphilicity of the AH attenuates protein-induced tubulation both *in vitro* and *in vivo*, whereas a super-amphiphilic AH variant (Arf1-F5W) enhances tubulation under both conditions^26^. AH insertions constitute a common mechanism of both membrane curvature generation and anchorage of a coat^6^, supported in the Arf1-only tubules by favorable electrostatic interactions between the positively charged inner surface of the G-domain scaffold and the negatively charged headgroups of acidic lipids in the membrane tubule outer leaflet (Fig. 4c). The membrane bilayer in the Arf1 tubules appears substantially thinner than in the MD simulations^32^ (Supplementary Fig. 7b-d).

The tubules are polar, with an A- and a B-end (Fig. 3). The regular arrangements observed in three distinct tubular lattices (corresponding to two-, three- and four-start helices with about 2× 16, 3× 18 and 4× 21 monomers per turn of 195 Å, 215 Å and 240 Å diameter tubules, respectively) resemble a close-packed rolled-up two-dimensional crystal, with dominant interactions along each helical strand and somewhat weaker inter-strand interactions. The similarity of the power spectra across 2D class averages (Supplementary Fig. 5) suggests that the broad range of tubular diameters (180-400 Å) share a similar local assembly of the coat. We therefore infer that the packing arrangement of the Arf1 scaffold allows generation of polymorphic membrane tubules of variable helical symmetry with diverse diameters and therefore curvatures. Similar observations have been made for other membrane tubules possessing a protein coat such as endophilin and ESCRT-III, which can support varying curvatures by a fractional addition or removal of endophilin dimers^45,46^ or CHMP2A-CHMP3 heterodimers^47^, respectively. For ESCRT-III, such short-range inter-filament interactions are thought to enable filament sliding, so that sequential removal (addition) of subunits could result in a stepwise decrease (increase) in tube diameter^47^. Modulation of short-range inter-filament interactions within a conserved overall lattice structure has also been attributed to variations in SNX1-coated membrane tubule diameter^48^, suggesting that membrane-templated modular assembly of repeating and consistent intermolecular interactions may be an underlying principle for the formation of tubules with variable diameters.

The crystalline G-domains in the reconstructed Arf1-GTPγS tubules were obtained from GUVs using high protein and nucleotide concentrations with prolonged incubation times. The densely packed Arf1 lattices appear to be relatively rigid; sparsely distributed Arf1 molecules would be expected to produce the soft meandering Arf1-tubes described previously^5,24^, some of which have such large diameters that their tubular nature could be detected using light microscopy^5,24^. Tubulation on GUV membranes is not observed at sub-micromolar Arf1-GTPγS concentrations, although Arf1 is still capable of recruiting other proteins^49^. At low concentrations, the degrees of freedom of membrane-bound activated Arf1 are restricted to lateral in-plane diffusion, rotational diffusion around the membrane normal and limited movement of the G-domain connected to the AH by the peptide linker, as suggested by MD simulations^32^ (Supplementary Fig. 7a).

We postulate that in our *in vitro* system, activated membrane-bound Arf1 molecules can self-associate reversibly to form two-dimensional crystals, supported by isolated low-resolution images that suggest the formation of such 2D Arf1-GTPγS lattices on planar membrane patches and on tubules (Supplementary Figs. 8c, 12c). As in other nucleated self-assembly processes (e.g. crystallization or fibrillation), incorporation of further membrane-bound Arf1-GTPγS molecules into the lattice becomes energetically favorable once a critical nucleus size is achieved. Formation of the crystalline array, generated by formation of the lateral contacts in interfaces 1 to 3, restricts the conformational space of the G-domains, pulling them to a constant distance of approximately 9 Å (Fig. 3c, g, Supplementary Fig. 7c). The inherent curvature of the G-domain protein lattice would result in a concomitant curvature of the supporting membrane, generating a protrusion (Fig. 7, Supplementary Movie 1a, b). At a point where the scaffold-lattice-generating forces outweigh those maintaining a flat bilayer, a membrane tubule can form with a diameter governed by the number of monomers completing the initial strand. This in turn can act as a template for the incorporation of subsequent monomers, resulting in the observed uniform Arf1-GTPγS membrane tubules.

**Fig. 7:**
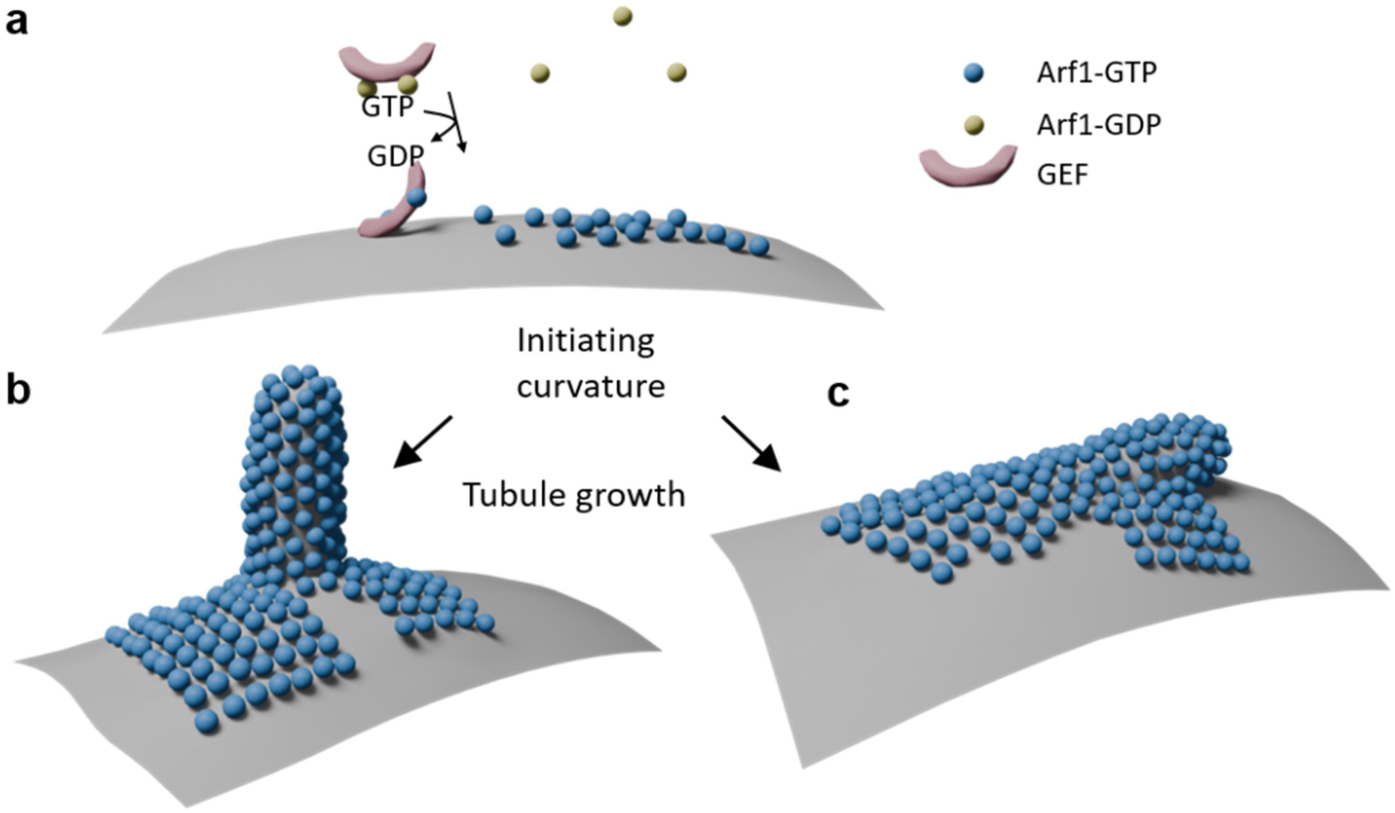
Proposed mechanism for Arf1-mediated membrane remodeling. **a** Membrane binding: In its GDP-bound state, Arf1 is monomeric and diffuses freely in the cytosol. Guanine nucleotide-exchange factors (GEFs, pale pink) recruit GDP-bound Arf1 (yellow) and catalyze GDP-to-GTP exchange, directing it to the membrane. In the GTP-bound state (blue), formation and exposure of the previously unstructured N-terminal amphipathic helix of Arf1 results in insertion into the outer membrane layer, initiating membrane curvature. **b, c** Oligomerization and curvature induction: At high local concentrations on the membrane, Arf1 can cluster to further enhance membrane curvature. Following formation of a sufficiently large nucleus, tubules may grow perpendicularly (**b**) or at an inclined angle (**c**) as observed occasionally in the cryo-EM micrographs (Supplementary Fig. 12).

Long, micrometer-scale tubule segments typically maintain a constant diameter (Fig. 2, Supplementary Figs. 2, 12), suggesting that initiation of the helical scaffold governs tubule dimensions. Nevertheless, both abrupt and gradual changes in diameter along a tubule are seen occasionally (Supplementary Fig. 12e-i), which may be attributed to the formation and propagation of tubule coat lattice defects. Also present in the electron micrographs are tubules as thin as 150 Å that in some cases appear to originate from/ fuse to wider tubules (Supplementary Fig. 2). If one assumes the same dimensions of the protein coat as in the wider tubes (compare Fig. 3, Supplementary Fig. 7d), then these thin tubules probably do not possess an inner lumen or exhibit a micellar character. It has been suggested that narrow Arf1 tubules may play a role in neck compression prior to vesicle scission^30^, so that such narrow lumen-less tubules may serve as a model for late stages of the fission process.

The conserved Arf1 lattice arrangement yields a packing area of approximately 1300 Å^2^ per monomer (around 76,000 molecule per μm^2^), which is of the same order of magnitude to that determined for Arf1 tubules in cells by fluorescence^27^ (taking STED resolution limits into consideration) and electron microscopy^28^. *In vivo*, the local concentration of activated Arf1 will at least in part be governed by the interplay of GEF activation (through GDP-GTP exchange) and GAP deactivation (through GTP hydrolysis), which is likely to result in a highly dynamic process. Arf1 tubule nucleation may occur where GEF activity is high and/or GAP activity is low, in keeping with data suggesting that this process is spatially and temporally regulated by GEFs^50^. In the structure of yeast GEF Gea2 in complex with the Arf-like protein Arl1 (where the membrane-binding, heel-like motif of Gea2 juxtaposes the membrane-binding face of Arf1), interface 1 would be occupied by the GEF and interface 1ʾ would be exposed in an orientation that could allow direct incorporation into the developing lattice at the A-end of a nascent helix (Supplementary Fig. 13a-c). As the tubes are polar, this would have the effect of transporting membranes unidirectionally from a ‘source’ membrane in which Arf1-GTP is produced.

As the nucleotide binding site is exposed on the outer surface of the tubules, it is accessible to GAPs; indeed, superposition of ArfGAP2^39^ to an Arf1 monomer in the tubules indicates that the GAP can bind with only minor clashes (Supplementary Fig. 13d-f). GAP action on Arf1-GTP, which stimulates GTP hydrolysis, would have three synergetic effects: (i) reorganization of the switch I and II regions to adopt the Arf1-GDP conformation, which is incompatible with the lattice structure; (ii) loss of Arf1-GDP contacts to the membrane tubule through dissociation of the AH from the membrane, which in turn (iii) would result in unfavorable lipid packing that has been suggested to drive fission^51^. Such disturbances of the lattice would result in point defects that could either lead to tubes of a different diameter as outlined above, or complete disruption of the scaffold to form a new (vesicular) membrane surface. *In vivo*, GTP hydrolysis and packing defect generation could be enhanced locally by ArfGAP1-binding to packing defects via its lipid packing sensor^52,53^.

In contrast to the tubule diameters of 150 - 400 Å observed here, Arf1-positive tubules in cells have been reported to be around 1100 Å in diameter using STED-based fluorescence microscopy (∼400 Å resolution)^27,54^, although a range of 170 - 1720 Å has been determined using FIB-SEM^28^. Thus, the lower limit of this intracellular Arf1 tubular diameter range closely matches that observed in our reconstituted Arf1 tubules, yet the higher end is well in excess. Whereas the *in vitro* tubular structures are maintained by G-domain contacts within the scaffold, the AH domains are free to undergo 2D rotational and translational lateral diffusion in the membrane, owing to conformational flexibility of the F13-K16 peptide linker that connects the AH to the G-domain scaffold. Such linker-mediated fluidity can locally create dynamic spaces both within the outer leaflet of the cylindrical membrane and between the membrane and the protein scaffold (Fig. 3c, g), which could allow accommodation of transmembrane cargo proteins. Larger luminal cargo protein domains could in turn lead to and stabilize larger diameter tubes similar to those observed in cells, with the Arf1 scaffold adopting lower curvatures.

Arf1 GDP/GTP state cycling triggers not only membrane recruitment, but also subsequent adaptor protein binding to form Arf1 heterocomplexes. The close packing of the Arf1 G-domains within the present tubules is in contrast to the loose packing observed for Arf1 in heterocomplexes^2,39,55–57^ (Supplementary Fig. 14), with little or no correspondence with the interfaces observed here. In COPI vesicles, Arf1 is found in a triangular arrangement^39^ (Supplementary Fig. 14a, b). AP-1:Arf1 tubules^56^ and AP-3:Arf1 heterocomplexes^55^ reveal Arf1 dimers at the membrane-proximal layer (Supplementary Fig. 14c-h). The face-to-face arrangement (correlating to interface 2b-2b) of Arf1 dimers in the AP-3:Arf1 heterocomplex may be a result of the heterocomplex formation in conjunction with the shape of the nanodisc edge (Supplementary Fig. 14f-h). While Arf1 dimerization has been discussed in previous studies^5,24,30^, our results indicate that membrane remodeling by Arf1 alone is driven primarily by myriad cooperative interactions, leading to a highly ordered directional oligomerization of Arf1. This Arf1 assembly can act as a well concentrated reservoir of membrane-bound Arf1-GTP for coatomer and adaptor proteins to recruit, an essential primary basis for directive intracellular membrane trafficking^50^.

A structure of membrane-bound mammalian Arf6 tubules has recently been presented^44^. Arf6, a paralogue of Arf1 that diverges most from Arf1 within the Arf family^58^, exhibits 68% sequence identity to Arf1 in mammalian organisms. In contrast to the polar Arf1 tubules presented here, with one monomer as a repeating unit, the Arf6 tubule is two-fold symmetric, with a repeating unit of four monomers that are related by two-fold C2 symmetry perpendicular to the helix axis (Supplementary Fig. 15d-f). Although a similar packing density of the G-domain is found and comparable interaction energies are achieved (Supplementary Table 3), none of the Arf6 interfaces correspond to those within the Arf1 helical assemblies. Despite several of the involved residues matching due to common binding motifs of the Arf proteins, it seems unlikely that Arf1 can adopt the Arf6 two-fold symmetric organization due to repulsive charge interactions within the tetrameric building block (Supplementary Fig. 16). On the other hand, there are no obvious indications that Arf6 cannot adopt the polar Arf1 tubule arrangement. Arf1 mutational analyses based on the Arf6 structure^44^ are also consistent with the polar Arf1 tubule structures presented here. The different lattice structures adopted by Arf1 and Arf6 might relate to functional differences between the two paralogues: Arf1 is localized predominantly to Golgi compartments, consistent with its role in secretory transport pathways^59,60^, whereas Arf6 is found mainly at the plasma membrane and on endosomal compartments, where it is involved in endocytic trafficking. Although the difference in tubule architecture could be a result of other factors, it is tempting to speculate that the unidirectionality of the Arf1 tubules compared to the bidirectional Arf6 tubules may reflect different functions of the two proteins.

The structural data presented here for the Arf1 tubules support the prominent role of Arf1 in membrane remodeling processes. Providing a high local concentration of GTP-loaded Arf1 in a lattice-like membrane-containing tubular assembly might serve as a reservoir for subsequent intracellular transport processes. The ability of Arf1 to remodel membranes through multiple organizational modes on membranes facilitates the multifaceted functions to which Arf1 has been attributed.

## Data availability

- Structural data supporting this study have been deposited in the Electron Microscopy Data Bank (EMDB) and Protein Data Bank (PDB) under the following accession numbers: EMD-19731, EMD-19732, EMD-19733 and PDB: 8S5C, 8S5D, 8S5E. Raw image data have been deposited to the Electron Microscopy Public Image Archive (EMPIAR)^61^ under accession number EMPIAR-12215.
- This paper does not report original computational code.
- Any additional information required to reproduce or further analyze the data presented in this study is available upon request from the corresponding authors.

## Quantification and statistical analysis

- Data collection and refinement statistics, along with the specific PDB and EMDB entries are provided in Supplementary Tables 1, 5.
- Standard deviations (1 σ) are reported in Supplementary Tables 2, 3 indicating the statistical variation of each metric.
- FSC measurements were used to estimate the resolution of cryo-EM maps, implemented in cryoSPARC^62^. The gold-standard FSC approach was applied, where FSC is computed from independently refined half-reconstructions. Detailed descriptions of this analysis are provided in the Results section, Figure legends, and Methods section.

## Acknowledgements

We thank Claudia Mueller, Anna Schoenberg and Fotis Kyrilis for technical assistance, Christian Tüting for data migrations and IT-related tasks, Andrea Scrima, Annette Meister, Christoph Parthier, and Fotis Kyrilis for helpful discussions. We are especially grateful to Pavel Afonine, Oleg Sobolev and Tom Terwilliger from the Phenix support team as well as Tristan Croll, the ISOLDE developer, Mikel Iceta Tena and Pablo Conesa from the Scipion team, and Helmut Gluender from ImageJ dealing with our particular requests. We also thank Johanna Hohgardt for preparing myr-Arf1-Alexa Fluor 488. Cryo-EM infrastructure (200 kV instrument) was supported by grants to P.L.K. The authors acknowledge the funding from the Federal Ministry for Education and Research (BMBF, 03Z22HN23 (to P.L.K.), 03Z2HN22 (to K.B.), and BMBF, 03Z22HI2 and 03COV04 (to M.T.S., K.B. and P.L.K.)), from the European Union through funding from the Horizon Europe ERA Chair “hot4cryo” project number 101086665 (to P.L.K.), and from the German Research Council (DFG, project-ID 436494874 – RTG 2670 (to K.B. and S.N.), INST 271/391-1 FUGG (to K.B.), and project-ID 391498659 – RTG 2467 (to M.T.S. and P.L.K.)). This work was also supported by FCT – Fundação para a Ciência e a Tecnologia, I.P., through MOSTMICRO-ITQB R&D Unit (DOI 10.54499/UID/04612/2025, UID/PRR/4612/2025) and LS4FUTURE Associated Laboratory (DOI 10.54499/LA/P/0087/2020).

## Author contributions

C.H., S.D. and S.N. prepared the protein, C.H. performed extensive cryo-EM sample preparation and optimization, F.H. and C.H. performed cryo-EM data acquisition, C.H., D.A.S. and A.D. carried out data analysis, M.T.S. assisted in data interpretation, C.H., S.D., S.N. conducted the biochemical experiments, P.L.K. performed the docking calculations, C.H. drafted the manuscript, assisted by D.A.S., M.T.S. and K.B.; K.B., C.H., D.A.S. and P.L.K. conceptualized the project, K.B., M.T.S. and P.L.K. provided input in various aspects of the work and supervised the project. All authors contributed to the final version of the manuscript.

## Additional information

Supplementary Information. The online version contains supplementary material available at …

Correspondence

Correspondence and requests for materials may be directed to and will be fulfilled by the lead contact, Kirsten Bacia upon request after completion of a Materials Transfer Agreement. This study did not generate new unique reagents.

## Methods

### Protein expression and purification

Myristoylated Arf1 (myr-Arf1) from *Saccharomyces cerevisiae* (UniProt P11076) was produced in a bacterial expression system according to protocols from Randazzo *et al.*^63,64^ and Zhang *et al.*^32^. Briefly, Arf1 (pOW12_ARF1) and N-myristoyl transferase (NMT1, UniProt P14743; pBB121_NMT1) were co-expressed in *E. coli* BL21 (DE3) in ampicillin and kanamycin supplemented LB medium to yield myr-Arf1 protein, which was purified by anion exchange chromatography (DEAE-Sepharose) followed by hydrophobic interaction chromatography (Phenyl Sepharose) and size exclusion chromatography (Sephacryl S100 16/600 HR). Labeling of myr-Arf1 with Alexa Fluor 488 maleimide (Invitrogen, Karlsruhe, Germany) was performed as described for Sar1p-CSSC^65^. Point mutations of selected residues were introduced by conventional site-directed mutagenesis methods using the pET24b(+) plasmid with DNA encoding for ARF1 (primers are listed in Supplementary Table 4). Arf1 mutants were co-expressed with NMT1 (pCDFDuet-1_NMT1) in LB medium containing kanamycin and spectinomycin, and purified as described for the wild-type protein.

### Giant unilamellar vesicles (GUVs)

All phospholipids were purchased from Avanti Polar Lipids (Alabaster, AL, USA) and ergosterol was purchased from Cayman Chemical (Ann Arbor, MI, USA). Fluorescent dyes were purchased from ThermoFisher Scientific (Waltham, MA, USA). GUVs were prepared from the major minor mix (MMM) of lipids according to Matsuoka *et al.*^66^ containing 34.4 mol% 1,2-dioleoyl-*sn*-glycero-3-phosphocholine (DOPC); 14.8 mol% 1,2-dioleoyl-*sn*-glycero-3-phosphoethanolamine (DOPE); 3.4 mol% 1,2-dioleoyl-*sn*-glycero-3-phosphate (DOPA); 5.4 mol% 1,2-dioleoyl-*sn*-glycero-3-phosphoserine (DOPS); 5.4 mol% L-α-phosphatidylinositol (soy-PI); 1.5 mol% L-α-phosphatidylinositol-4-phosphate (PI(4)P, from porcine brain); 0.5 mol% L-α-phosphatidylinositol-4,5-bisphosphate (PI(4,5)P2, from porcine brain); 1.3 mol% 1,2-dioleoyl-*sn*-glycero-3-(cytidine diphosphate) (CDP-DAG) and 33 mol% ergosterol. Lipids were dissolved in 2:1 (v/v) chloroform:methanol and, when indicated, supplemented either with 0.1 mol% of the fluorescent lipid analogue 1,1’-dioctadecyl-3,3,3’,3’-tetramethylindocarbocyanine perchlorate (DiI-C_18_) or with 0.1 mol% of the fluorescently labelled lipid 1,2-palmitoyl-*sn*-glycero-3-phosphoethanolamine-(DPPE)-Atto647N at a total lipid concentration of 10 mg ml^-1^ and spread on two preheated indium-tin-oxide (ITO) coated glass slides (Delta Technologies, Loveland, CO, USA) at 65 °C to obtain a thin lipid layer after evaporation of the solvent. The two glass slides were assembled with a 1.2 mm thick silicon ring to form an electroformation chamber, filled with an aqueous sucrose solution (500 mOsmol kg^-1^). Electroformation^67^ was performed using a 1.3 V (pp), 10 Hz sinusoidal voltage for 7 h. GUVs were harvested by overnight incubation in 500 mOsmol kg^-1^ glucose at 4 °C.

### Arf1-mediated tubulation

Tubulation was performed in HKM-buffer (20 mM 4-(2-hydroxyethyl)-1-piperazineethanesulfonic acid (HEPES), pH 6.8, 50 mM potassium acetate, 1.2 mM MgCl_2_) by incubating 0.3 μl GUVs with 10 μM myr-Arf1, 1 mM guanosine 5’-O-(3’-thiotriphosphate) (GTPγS) and 2.5 mM ethylenediaminetetra-acetic acid (EDTA) in a final volume of 10 μl at 22 °C. Mutant myr-Arf1 proteins were treated analogously. To analyze the co-localization of Arf1 with membranes by confocal fluorescence microscopy, 6 μM partially fluorescent-labelled myr-Arf1 (40% myr-Arf1-Alexa Fluor 488) was used instead. For confocal microscopy, 3 μl of the tubulation reaction was immediately transferred to 2%-gelatin-coated glass coverslips equipped with double-sided adhesive tape and covered by a second glass coverslip to form an enclosed microscopy chamber. Otherwise, incubation was performed in the reaction tube for the indicated incubation times.

### Confocal microscopy

Arf1 tubulation was imaged on an inverted Zeiss Axio Observer 7 microscope (Carl Zeiss Microimaging, Jena, Germany), equipped either with a laser scanning system (LSM 980) or a CSU-X1 spinning disk module (Yokogawa Electric, Musashino, Japan) using a C-Apochromat 40 ×, NA 1.2 water immersion objective. Excitation of the dyes Alexa Fluor 488, DiI-C_18_ and Atto647N was achieved by diode lasers at 488 nm, 561 nm and 639 nm, respectively; fluorescence emission was collected by the Airyscan2 detector (LSM 980) or by using a BP 525/50 and BP 640/132 dual-band emission filter (spinning disk microscope). The two-color images were acquired in the line-wise multi-track mode. Confocal microscopy under cryo conditions (-196 °C) was performed using an upright Zeiss Imager.Z2 microscope with a laser scanning system (LSM 900, Carl Zeiss Microimaging, Jena, Germany) equipped with the cryo-stage CMS196M (Linkam Scientific Instruments, Tadworth, UK). Vitrified grids were mounted onto the cryo-stage and visualized under nitrogen atmosphere maintained at -196 °C to prevent sample thawing. Excitation and emission settings were the same as described for the LSM 980. Using the Carl Zeiss ZEN software, a non-linear intensity look-up table with γ = 0.45 was applied where indicated.

### Cryo-EM grid preparation and data acquisition

For cryo-EM grid preparation, 3.5 μl of the tubulation reaction sample was applied to both sides of a Quantifoil holey carbon grid (R2/1, 200 mesh). The grids were pre-treated in a plasma cleaner (Harrick Plasma PDC-32G, Ithaka, NY, USA) for 25 seconds. Vitrification was performed using the Leica GP1 grid plunger (Leica Microsystems, Wetzlar, Germany) with 80% humidity, with back-blotting for 10 seconds followed by rapid freezing in liquid ethane. Data collection was carried out using a Glacios microscope (ThermoFisher Scientific, Waltham, MA, USA) operated at 200 keV. Micrograph movies were collected with a Falcon 4i direct electron detector (ThermoFisher Scientific, Waltham, MA, USA), at a magnification of 92,000 ×, yielding a pixel size of 1.53041 Å. The dose rate was set to 21 e^−^ per pixel per second, and a total dose of 50 e^−^ per Å^2^ was fractionated over 40 frames. The data collection process, including defocus values ranging from -0.6 μm to -2.8 μm, was managed using the EPU software (version 2.2.0.65REL). A total of 12,483 movies were acquired. Additional datasets for screening tubular and lattice shapes were collected with a Falcon 3 electron detector (ThermoFisher Scientific, Waltham, MA, USA) at the same magnification, resulting in a pixel size of 1.5678 Å.

The tubules were initially visualized and tube diameters were measured using ImageJ2 software (available at http://imagej.org)^68^. To register electron and fluorescence microscopy images, the ImageJ plugin Correlia^69^ was used.

### Image processing of Arf1 tubules

Cryo-EM image analysis was performed using cryoSPARC v4.3.1 and v4.4.2^62^ (Supplementary Fig. 3). A total of 12,483 raw movies were used for electron beam-induced motion correction using MotionCor2^70^. Corrected micrographs were then subjected to ‘patch-CTF estimation’. Particle picking was performed using the ‘filament tracer’ with a template derived from 2D classification of manually picked filaments (box size 384). Each micrograph was manually revised to assess and correct the template-picking results. Initially, 649,513 particles/segments were extracted with 75% overlap between consecutive segments. These were subjected to a first 2D classification to provide the 2D class averages for an estimation of the helical symmetry parameters. To determine a range of potential solutions of helical symmetry parameters, the sum of power spectra of the segments belonging to one 2D class average of the respective 2D class was calculated using the ‘Average Power Spectra’ routine in cryoSPARC v4.3.1. Subsequently, these averaged power spectra were analyzed by the web tool helixplorer (https://rico.ibs.fr/helixplorer). The helical rise of 4.3 Å approximated in this manner was used to adjust the separation distance between segments for the following extraction. 1 489,051 particles/segments were extracted with a final separation distance of 21.5 Å between individual segments, and then parsed using several rounds of reference-free 2D classification with precise visual inspection of diameter classes as well as power spectra to discriminate between the small differences in tube diameter. 2D class averages were then combined according to equal diameters and power spectra. Due to the broad distribution of filament diameters with only tiny differences, we focused on filament diameters from 185 to 215 Å, which comprised 1/3 of the dataset. A total of 485,140 particles/segments were selected after multiple rounds of 2D classification using cryoSPARC v4.3.1. Two homogenous subsets of data representing the two groups of a filament diameter of 195 Å and a filament diameter of 215 Å were selected for further processing and analysis steps using a dataset of 180,983 segments classified as tubules with a diameter of 195 Å and a dataset of 31,369 segments for tubules with a diameter of 215 Å. The most abundant class of 200 Å diameter tubules (Fig. 2b) suffered from severe inhomogeneities that could not be separated by extensive rounds of 2D classification and averaged power spectra generation and, therefore, was not used for further refinement. Each averaged power spectrum was analyzed in detail with the help of the web tool helixplorer to assess the final helical symmetry parameters for each filament diameter. Estimates for the pitch (first layer line with maximum intensity) as well as for the axial rise (layer line that merges with the meridian) were made according to the layer line positions and the corresponding intensities (Supplementary Fig. 4). Several potential symmetries for each of the two diameters were tested in a 3D reconstruction using the Iterative Helical Real Space Reconstruction (IHRSR) algorithm^71,72^ that is based on a single-particle approach while imposing and refining helical symmetry, implemented in the ‘Helical Refinement’ routine in cryoSPARC. Only symmetries that gave a reasonable density distribution were subjected to iterative 3D helical refinements to further refine the symmetry parameters one by one. The resulting maps were investigated for high-resolution features, such as secondary structures and aromatic side chains, as well as for the quality of the fit of the atomic model (PDB: 2ksq) to decide on map integrity. Multiple rounds of iterative 2D classifications, selections and 3D helical refinements were performed to produce the final helical map using cryoSPARC v4.4.2. After obtaining the final helical reconstruction, the datasets with the two diameters were corrected for reference-based motion (job type ‘Reference-based Motion Correction’) followed by a final 3D helical refinement with the previously found helical parameters (summarized in Supplementary Table 1). The best map for each diameter was subjected to 3D classification, and a single homogenous 3D class containing 127,727 and 27,285 segments for each diameter of 195 and 215 Å, respectively, was chosen for a final helical refinement.

Using a soft protein-only mask, the resolution of the maps was estimated at the Fourier shell correlation (FSC) cut-off criterion of 0.143^31^ to be 3.1 Å and 3.8 Å for the 195 and 215 Å diameter tubules, respectively.

Local resolution estimations on both cryo-EM maps (195 Å and 215 Å tubules) were performed using the postprocessing pipeline in cryoSPARC v4.4.2. Maps were sharpened with global B-factors of -108.5 (195 Å tubules) and -74.2 (215 Å tubules). Additional low-pass filtering according to the local resolution estimates of the final maps imported into RELION was performed in RELION^73,74^ to implement independent sharpening and filtering for the protein and for the membrane to be shown together as an assembled map. Appropriate local resolution filtering to better render a single map at a reasonable threshold was not achievable. Maps for local resolution filtering were sharpened with B-factors of -78.0 (195 Å tubules) and -39.2 (215 Å tubules) in RELION to visualize the membrane. Figures were generated in ChimeraX^75,76^ and Chimera^77^.

### Model building and refinement

Individual Arf1 molecules were clearly distinguishable in the cryo-EM 3D map of the Arf1-coated tubules at 3.1 Å resolution (195 Å tubules). The structure of the G-domain of GTP-bound yeast Arf1 from the solution NMR structure (PDB: 2ksq^11^; omitting the myristoyl chain, the N-terminal residues 1-15 and the C-terminal residues 179-181) and the well resolved nucleotide GTPγS (GSP) could be readily fitted using Coot (version 0.9.7 EL)^78^. In order to represent all possible intermolecular interactions along the tubules during the structure refinement, a cluster of seven subunits (one central protein molecule (chain A) with six surrounding molecules generated by helical symmetry) was extracted (Fig. 3d) for real-space refinement in Phenix 1.21.5187^79^, applying strict non-crystallographic symmetry (NCS) constraints throughout. The atomic model was subjected to interactive all-atom molecular dynamics flexible fitting using ISOLDE 1.6 (Interactive Structure Optimization by Local Direct Exploration)^80^ to optimize the initial fit to the density map before entering the Phenix real-space refinement pipeline. Ligand restraints for GSP and the magnesium ion were generated using the ‘electronic Ligand Builder and Optimization Workbench’ (elBOW)^81^, integrated in the Phenix suite. Several cycles of real-space refinement were performed in Phenix (phenix.real-space-refine^82^) incorporating simulated annealing protocols, strict NCS constraints, secondary structure restraints, Ramachandran restraints, and ligand restraints. Between refinement cycles, manual adjustments were made in Coot followed by further Phenix real-space refinement. The final model consists of the Arf1 asymmetric unit (chain A); in the pdb deposition (8s5e) the six neighboring symmetry related molecules are included. Validation and statistics for the refined model were obtained using MolProbity^83^ and Phenix validation tools, as summarized in Supplementary Table 5. Model quality was evaluated using map-to-model correlation coefficients, geometry indicators, and Fourier shell correlation (FSC) analysis between the map and the model. All molecular graphics were prepared with ChimeraX^75,76^ or Coot^78^.

### Energy calculations of the interfaces

High Ambiguity Data-driven biomolecular DOCKing (HADDOCK) runs were performed to analyze the energetics of the interfaces between individual Arf1 molecules within the tubular arrangement using the HADDOCK web server at https://alcazar.science.uu.nl/ (version 2.4)^35^. The applied refinement procedure only included the water refinement stage of the HADDOCK protocol while skipping the docking step to perform a qualitative correlation of the calculated energetics with binding affinities for transient protein-protein interactions^37^. The protocol consisted of a short molecular dynamics (MD) simulation using an explicit solvent shell, i.e. a thin and optimized water layer surrounding each protein. Settings were as follows: (1) solvation of the dimeric Arf1-Arf1 complexes in an 8 Å shell of TIP3P water by (i) 40 energy minimization steps with the protein fixed (Powell minimiser) and (ii) 2 × 40 energy minimization steps with harmonic position restraints of the protein (*k* = 20 kcal · mol^-1^Å^-2^); (2) final water refinement with a gentle simulated annealing protocol using MD in Cartesian space: (i) heating period: 500 MD steps at 100, 200 and 300 K while applying position restraints (*k* = 5 kcal · mol^-1^Å^-2^) on the protein except for backbone and side chains at the interface; (ii) sampling stage: 1250 MD steps while applying weak (*k* = 1 kcal · mol^-1^Å^-2^) position restraints on the protein except for backbone and side chains at the interface, and (iii) cooling stage: 500 MD steps at 300, 200 and 100 K while applying weak (*k* = 1 kcal · mol^-1^Å^-2^) position restraints on the protein except for backbone and side chains at the interface. The temperature was kept constant by weak coupling to a reference temperature bath using the Berendsen thermostat^84^. The equations of motion were integrated by using a time step of 2 fs. Non-bonded interactions were taken into account using the OPLS force field^85^ (cut-off 8.5 Å). The electrostatic potential was calculated by using a shift function while a switching function (between 6.5 and 8.5 Å) was used to define the van der Waals potential. A total of 20 structures were generated for each interface.

### Flotation Assays

For liposome binding reactions, 4 μM Arf1 or respective variants were incubated in a total volume of 80 μl with vesicles (0.5 mM lipid) of the same lipid mixture as used for the GUVs (labelled with 2 mol% DiI-C_18_) and 0.1 mM nucleotide. The reaction buffer contained 20 mM HEPES pH 7.4, 1.2 mM MgCl_2_, 50 mM NaCl, 15 mM potassium acetate and 2.5 mM EDTA to facilitate nucleotide exchange. After incubation for 60 min at 30 °C, 2 μl of a MgCl_2_ solution were added to a final concentration of 2 mM Mg^2+^ in order to stabilize the formed complex. The reaction was mixed with 50 μl of a 2.5 M sucrose solution in HKM-buffer. 110 μl of the mixture were transferred to a centrifugation tube and overlaid with 100 μl 0.75 M sucrose in HKM-buffer, and finally with 20 μl HKM-buffer as the top layer. Samples were placed in a TLA-100 rotor (Beckman Coulter, Krefeld, Germany) and centrifuged for 25 min at 24 °C and 100,000 rpm in an Optima-XP ultracentrifuge (Beckman Coulter, Krefeld, Germany) with acceleration and deceleration rates set to 5 (2.5 minutes to maximum speed) and 7 (6 minutes to rest), respectively. 50 μl of top and bottom layers were harvested to recover liposome-bound and unbound protein, respectively. The amount of recovered lipid was determined via the UV/Vis absorption of membrane-incorporated dye and samples brought to equal lipid concentrations by dilution with HKM-buffer. Proteins were assayed by 15% SDS-PAGE. Gels were loaded with top-and bottom fractions from flotation assays and stained with Coomassie brilliant blue.

