## Supplemental Material for "Structural basis of Arf1-driven membrane tubulation"

The document includes 5 tables and 16 figures. Additionally, one movie is provided.

### Supplementary Information

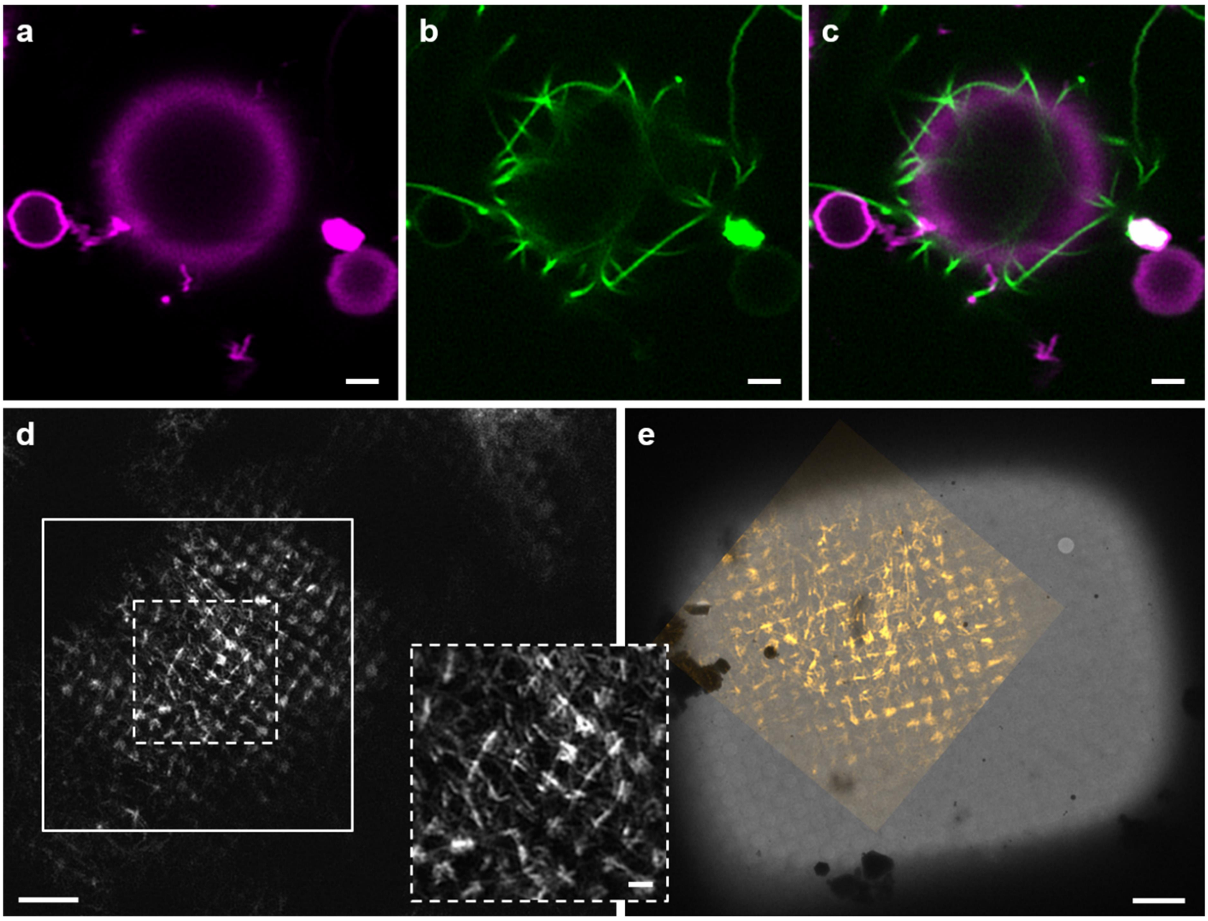


**Supplementary Figure 1**: Confocal microscopy studies on Arf1-mediated tubulation.

**a-c** Confocal microscopy images of DPPE-Atto647N fluorescently labeled membranes excited at 639 nm (magenta, (**a**)) incubated with GTPγS and green fluorescently labeled myr-Arf1 excited at 488 nm (**b**) showing straight, rigid lipid tubules that detach from the vesicular membrane. The overlay image (**c**) demonstrates that Arf1 is localized on the membrane deformations. Scale bars: 2 µm

**d** Representative grid square of a cryo-fluorescence microscopy image of vitrified myr-Arf1-GTPγS tubules on a Quantifoil R2/1 grid. Membranes stained with the fluorescent lipid analogue dye DiI-$C_{18}$ incorporated in the bilayer were excited at 561 nm and the emission was collected using an Airyscan2 detector. The inset with dashed lines shows a zoomed-in image of the tubules demonstrating the uniform distribution of the tubules over the grid square. Positions of the holes in the Quantifoil film are clearly visible.

**e** Superposition of the cryo-fluorescence image from the inset marked by a solid line in (**d**) with the same grid square detected by cryo-electron microscopy. The overlay shows the presence of tubules in thicker ice regions of the grid square, which are not visible in the EM overview image. The correlation of the two images was performed by Correlia[1]. Scale bars: 10 μm, inset 2 μm

**a**

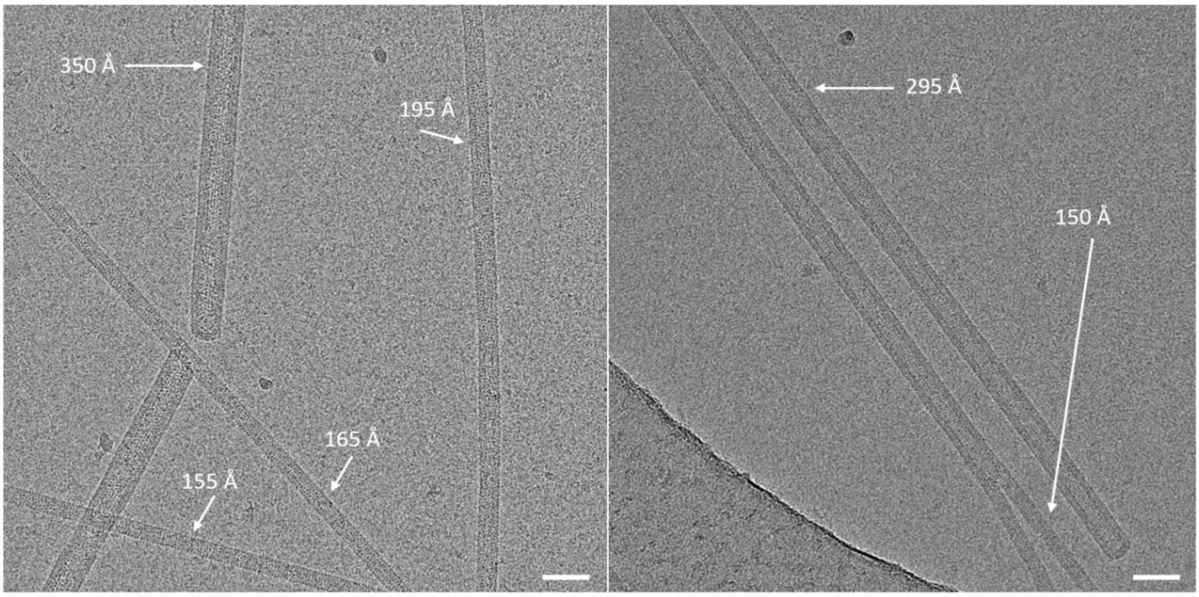


**b**

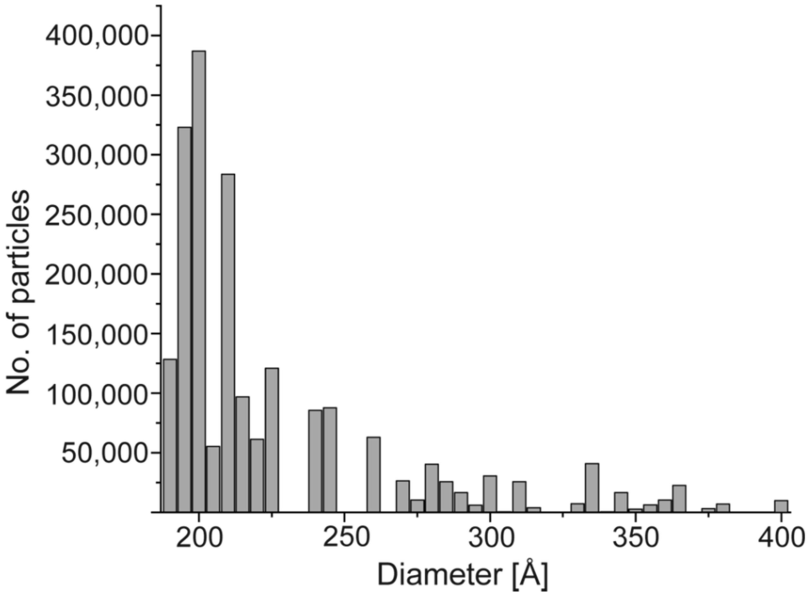


**Supplementary Figure 2**: Cryo-electron micrographs of the thinnest tubules observed. The respective diameters of the tubules are indicated (**a**).

**b** Size distribution plot of tubule diameters observed in the tubulation preparations. Only the most abundant diameters within a range of 180 to 280 Å (see Fig. 2b) were selected for further iterative 2D classification steps. Scale bars: 500 Å

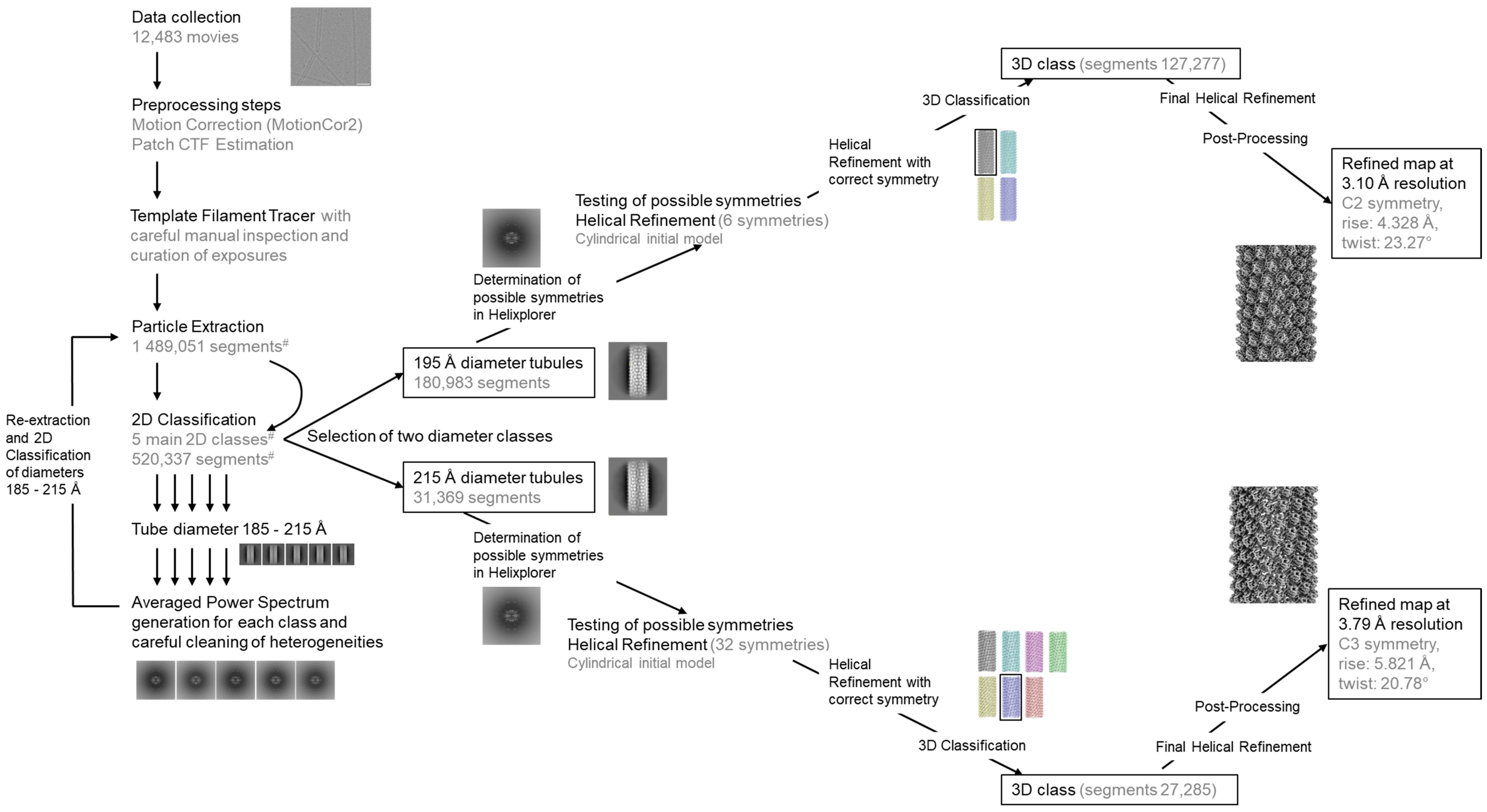

Data collection
12,483 movies
Preprocessing steps
Motion Correction (MotionCor2)
Patch CTF Estimation
Template Filament Tracer with careful manual inspection and curation of exposures
Particle Extraction
1 489,051 segments#
2D Classification
5 main 2D classes#
520,337 segments#
Tube diameter 185 - 215 Å
Averaged Power Spectrum generation for each class and careful cleaning of heterogeneities
Re-extraction and 2D Classification of diameters 185 - 215 Å
Selection of two diameter classes
195 Å diameter tubules
180,983 segments
215 Å diameter tubules
31,369 segments
Determination of possible symmetries in Helixplorer
Testing of possible symmetries
Helical Refinement (6 symmetries)
Cylindrical initial model
Helical Refinement with correct symmetry
3D Classification
3D class (segments 127,277)
Final Helical Refinement
Post-Processing
Refined map at 3.10 Å resolution
C2 symmetry,
rise: 4.328 Å,
twist: 23.27°
Determination of possible symmetries in Helixplorer
Testing of possible symmetries
Helical Refinement (32 symmetries)
Cylindrical initial model
Helical Refinement with correct symmetry
3D Classification
3D class (segments 27,285)
Final Helical Refinement
Post-Processing
Refined map at 3.79 Å resolution
C3 symmetry,
rise: 5.821 Å,
twist: 20.78°

**Supplementary Figure 3**: Cryo-EM data analysis workflow.

After several iterative 2D classification and particle extraction rounds, two homogenous datasets for the 195 Å diameter and the 215 Å diameter tubules were obtained and their helical symmetry was determined using Helixplorer (https://rico.ibs.fr/helixplorer). All processing and helical refinement steps were performed with cryoSPARC[2] according to the IHRSR[3,4] procedure. A detailed description of the individual processing steps is provided in the Methods section.

[#] Indicated values correspond to the pipeline after re-extraction and adjustment of the separation distance between segments with initial estimates for the helical parameters (see Methods section for more information). The initial 2D classification was carried out with 649,513 extracted particles based on a separation distance of 90 Å between consecutive segments (corresponding to 75% overlap of the helically arranged segments).

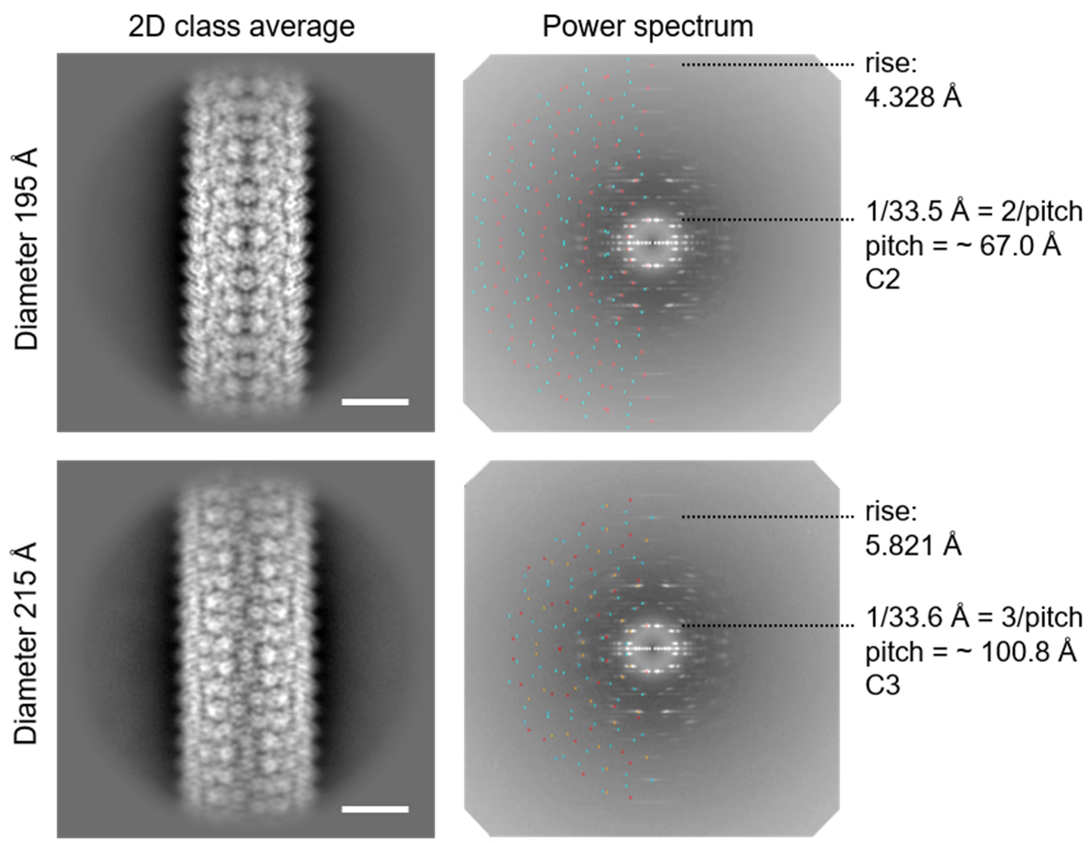


**Supplementary Figure 4**: Helical symmetry analyses.

2D class averages are shown for the two diameters 195 and 215 Å with their corresponding power spectra. The power spectra were created as the sum of the power spectra of the individual segments of the corresponding 2D class average using the cryoSPARC[2] suite. The layer lines indicating pitch and rise are highlighted. The calculated positions of the first maximum of each Bessel function from Helixplorer-1 (https://rico.ibs.fr/helixplorer/) are overlaid on the left half of the power spectrum. For clarity, expected positions of layer lines of higher Bessel orders were neglected.

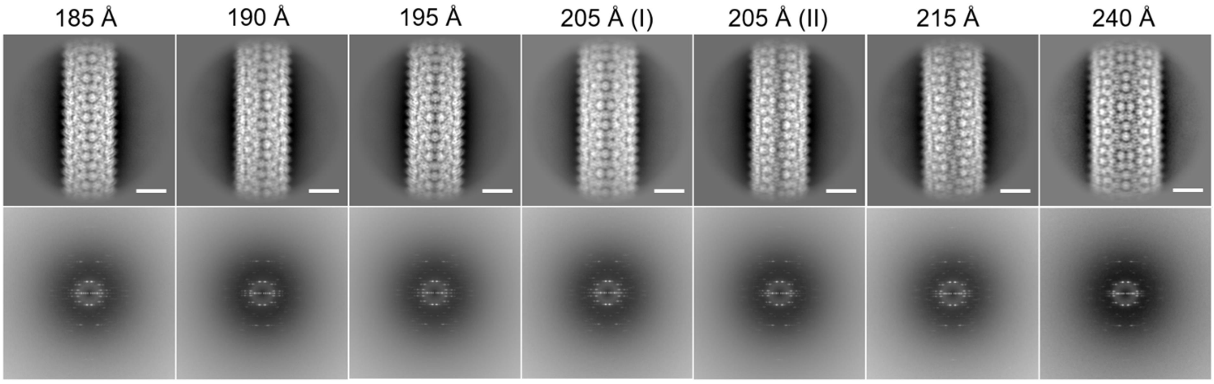


**Supplementary Figure 5**: Gallery of selected 2D class averages of tubules that differ in diameter (upper panels) and their respective power spectrum (lower panels) showing the consistency in the appearance of the power spectra.

The specific diameter is indicated at the top. Notably, the 205 Å tubules displayed two distinct power spectra, indicating the presence of two distinct classes. Power spectra were created as the sum of the power spectra of the individual segments of the corresponding 2D class average using the cryoSPARC[2] suite. Scale bars: 100 Å

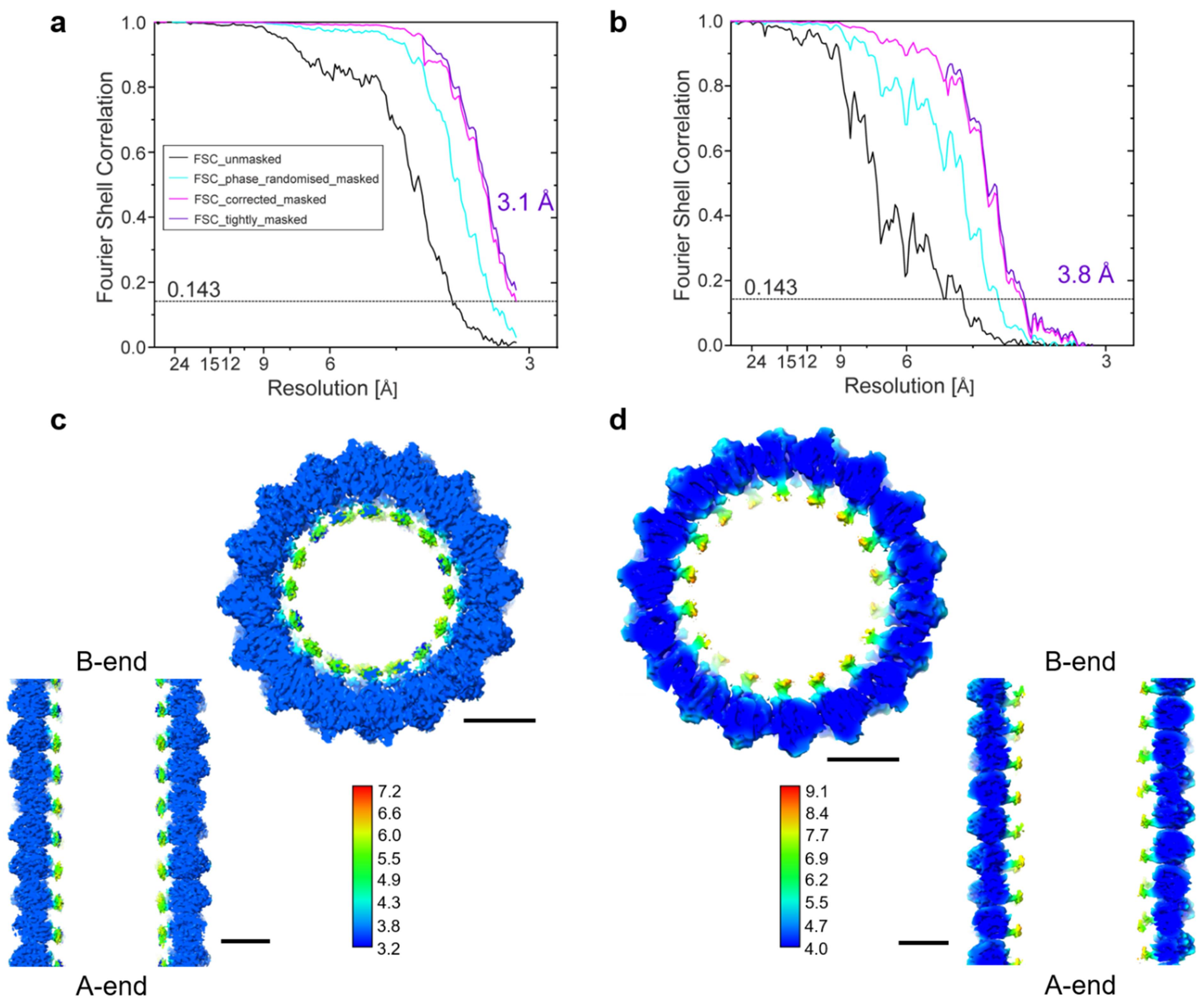


**Supplementary Figure 6**: Global and local resolution of the cryo-EM reconstructions of the two Arf1 tubules.

**a**, **b** Plots showing the Fourier shell correlation (FSC) versus spatial frequency for both diameters (195 Å (**a**), 215 Å (**b**)) after the post-processing step with FSC unmasked maps in black, the FSC phase randomized masked maps in cyan, the FSC corrected masked maps in magenta and the FSC tightly-masked maps in purple. The resolutions of the reconstructions were estimated to be 3.1 Å (195 Å) and 3.8 Å (215 Å) using the 0.143 cut-off criterion[5]. As the Nyquist limit is reached (pixel size 1.53041 Å), the FSC curve does not drop to zero after phase randomization (195 Å tubules (**a**)).

**c**, **d** Local resolution estimates mapped onto the cryo-EM density of the 195 Å (**c**) and 215 Å (**d**) diameter tubules in top-down view of central slices perpendicular to the helical axis and central slices in side view. Local resolution values in Å are shown by the color bar. Scale bars: 50 Å

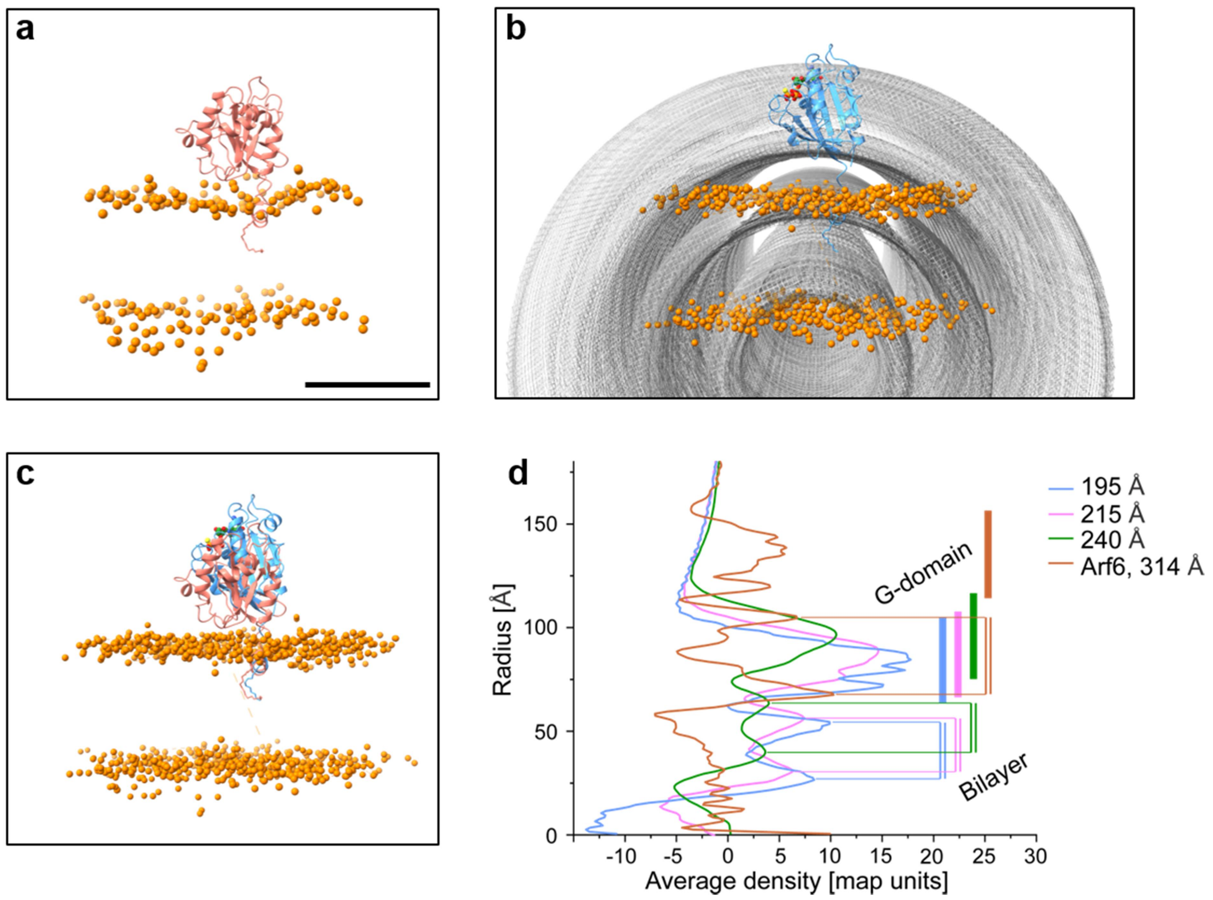


**Supplementary Figure 7**: Comparison of MD simulation results with the tubule structures.

**a** Representative snapshot of membrane-bound Arf1 from MD simulations[6]. Scale bar for **a**-**c**: 35 Å

**b** Cylindrically averaged density map of the 195 Å tubule (mesh representation) overlaid with Arf1-GTPγS (blue, ribbon representation with GTPγS shown as spheres). As the AH could not be resolved in the tubular structures, residues 2-15 including the N-terminal myristoyl chain were modelled according to MD trajectories. The flat phospholipid bilayer from MD simulations (phosphorus atoms shown as orange spheres) was positioned tangentially to the tubular membrane outer leaflet. Compared to the tubular bilayer, the flat bilayer appears to be significantly thicker.

**c** Superimposition of Arf1 in the phospholipid bilayer from MD simulations with Arf1 from the tubular structures. The G-domains of lattice-like tightly arranged Arf1 in the tubules appear to be much further away from the membrane than the G-domain of monomeric Arf1.

**d** Radial density profiles of Arf1-GTPγS tubules (blue, 195 Å; pink, 215 Å; green, 240 Å) and the Arf6 tubule (brown, 314 Å). G-domain and bilayer regions are indicated.

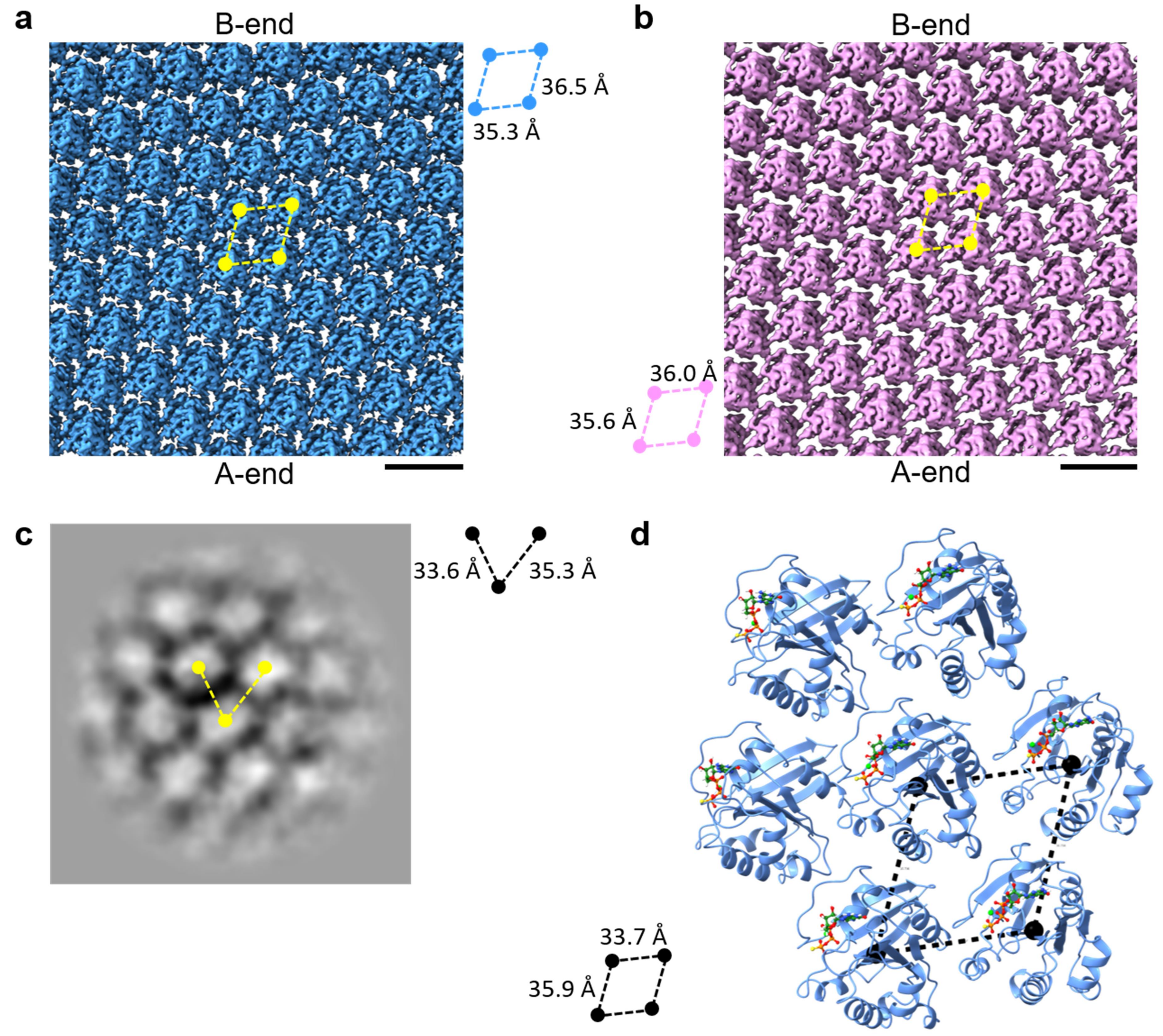


**Supplementary Figure 8:** Consistency of Arf1 2D lattices across tubular and planar projections.

**a, b** Cylindrical projections of unrolled 3D EM density maps for 195 Å (**a**) and 215 Å (**b**) diameter tubules. The intermolecular distances are indicated next to the respective map (blue: 195 Å, pink: 215 Å). Scale bars: 50 Å

**c** 2D class average of planar structures found in some of the electron micrographs (see Supplementary Fig. 12c), consistent with intermolecular distances observed in the tubules. Limited particle counts precluded 3D reconstruction.

**d** Refined structure of the Arf1 helical assembly (ribbon representation) from the 195 Å tubules, highlighting a central Arf1 molecule surrounded by six neighbors. The nucleotide is shown in stick representation, with the magnesium ion as a green sphere. Dashed black lines indicate inter-protein distances within the lattice.

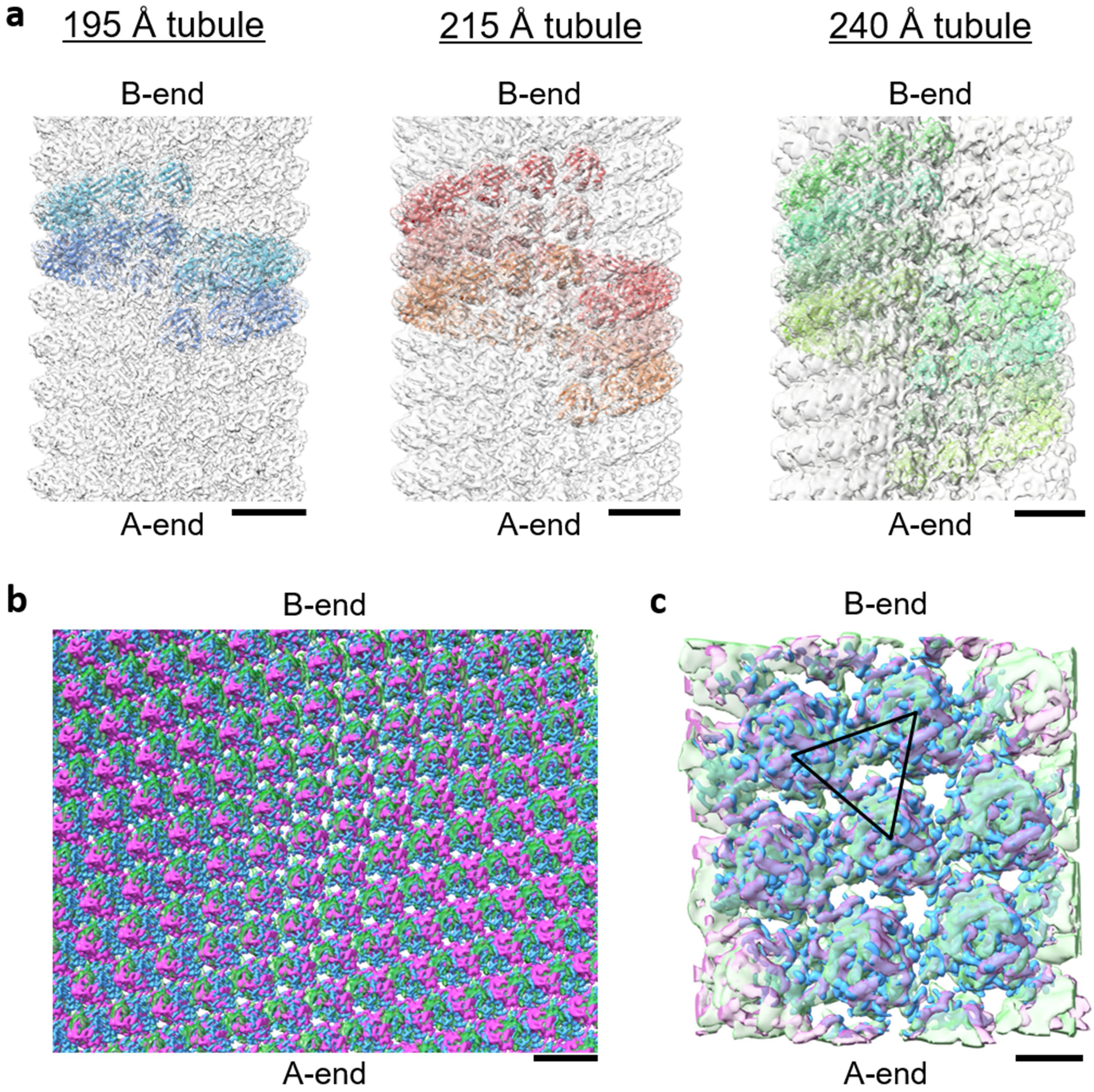


**Supplementary Figure 9**: Cryo-EM map of a 240 Å diameter tubule reveals a similar packing arrangement to 195 Å and 215 Å tubules.

**a** Side view of the high resolution cryo-EM maps of the 195 Å and 215 Å tubules and low resolution cryo-EM map of the 240 Å diameter tubule. Start and finish ends of the polar tubules are labeled with A-end and B-end, respectively. Individual filaments of the 2-start (195 Å), 3-start (215 Å) and 4-start (240 Å) helices are shown as cartoons in shades of blue, red/orange and green, respectively. Scale bars: 50 Å

**b** Overlay of the unrolled real-space maps for the 195 Å (blue), 215 Å (pink) and 240 Å (pale green) diameter tubules. Scale bar: 20 Å

**c** Zoomed-in superposition of three cryo-EM density maps demonstrates the conserved lattices among tubules of different diameters. The triangular arrangement for the three interfaces is indicated by black solid lines. Color-coding as before. Scale bar: 20 Å

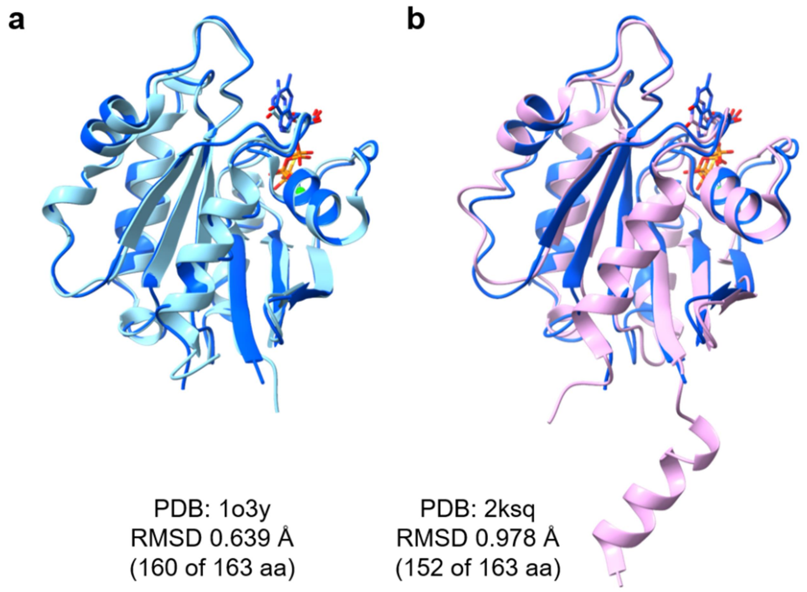


**Supplementary Figure 10**: Comparison of the Arf1-GTPγS model (dark blue) within the tubular arrangement to related structures of Arf1 in ribbon representation with corresponding global root-mean-square deviation (RMSD) values upon superimposing backbone $C_\alpha$ atoms. Nucleotides are shown in stick representation.

**a** Crystal structure of mouse Arf1[7] (cyan; PDB: 1o3y, resolution 1.5 Å).

**b** Solution NMR structure of yeast Arf1 in a bicelle-bound conformation[8] (pink; PDB: 2ksq.10).

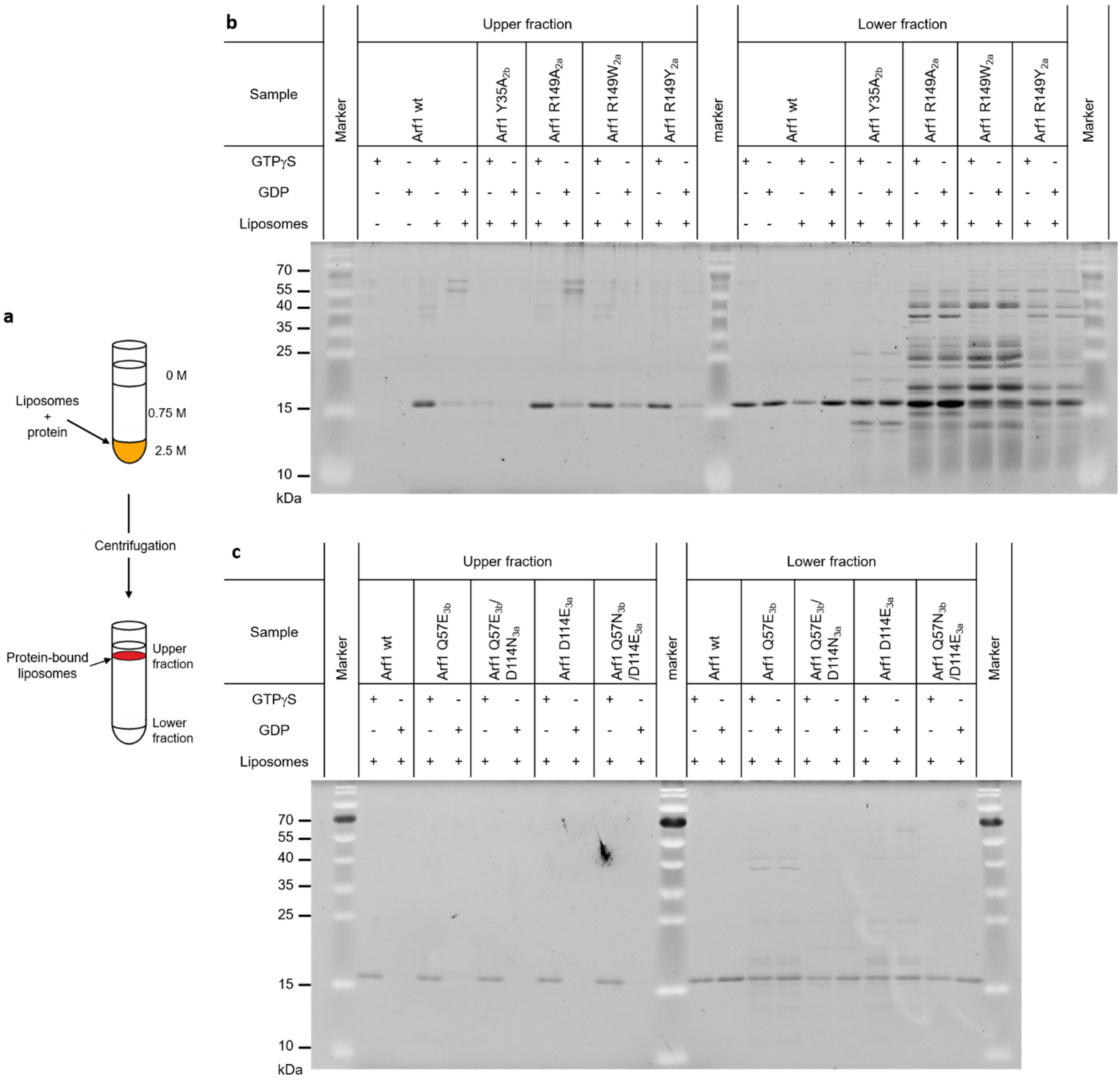


**Supplementary Figure 11**: Liposome flotation assay.

**a** Principle of the liposome flotation assay. Liposomes incubated with a putative membrane-binding protein are mixed with a highly concentrated sucrose solution (2.5 M) and overlaid with a less concentrated sucrose solution (0.75 M) and finally with buffer (0 M sucrose). Upon centrifugation, a continuous sucrose gradient is formed, in which (low density) liposomes will float near the top (with protein if bound), whereas (high density) non-bound protein collects at the bottom of the centrifugation tube.

**b, c** SDS-PAGE analyses of the liposome flotation assay of wild-type Arf1 and Arf1 proteins with mutations in interface 2 (**b**) and interface 3 (**c**) residues. Wt-Arf1 was either incubated with the nucleotide GTP$\gamma$S, which causes exposure of the N-terminal helix and binding to membranes, or GDP, which maintains the soluble state. A negative control was performed by omitting the liposomes. The Y35$A_{2b}$ mutant (interface 2, **b**) is deficient in membrane binding as almost no protein was observed in the upper, liposome-containing fraction. All other interface 2 mutants (**b**) are detected in the upper fraction, indicating their ability to bind to membranes. Protein preparations of the R149 mutants exhibited a number of impurities that do not bind membranes. The weak protein bands in the upper fraction observed in the GDP-bound samples of wt and R149$A_{2a}$ might arise from abundant membrane-binding proteins from the host organism *E. coli*. All interface 3 (**c**) mutants appear to bind to membranes as protein could be observed in the upper, liposome-containing fraction. As an excess of protein was added to the flotation reaction, all proteins were also detected in the lower fraction. Molecular weights of the marker bands (PageRuler) are listed on the left side of the gel in kDa.

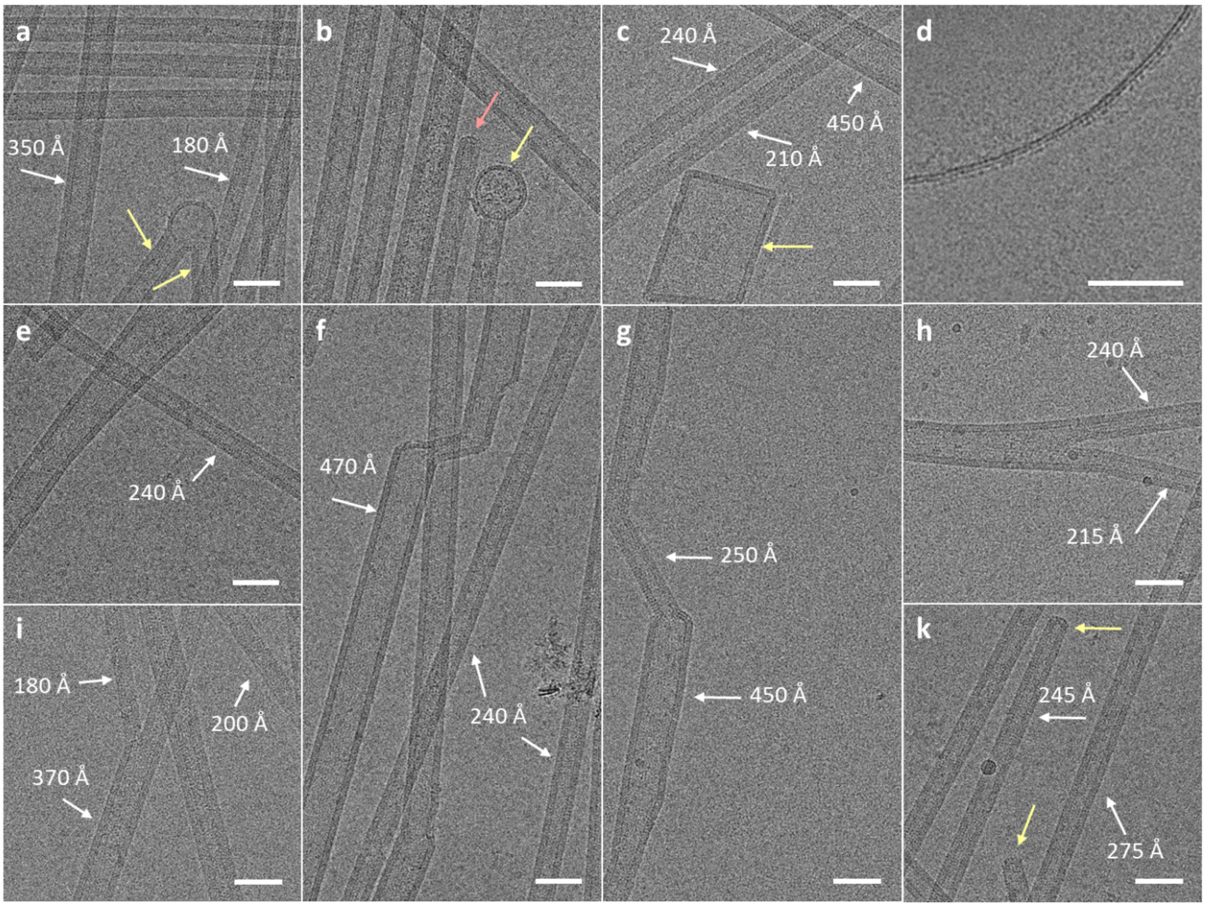


**Supplementary Figure 12**: Selection of cryo-electron micrographs that show membrane morphologies beyond long, rigid tubules. Examples of tubules with various diameters are indicated.

**a** Two tubules emanating from a single vesicle (yellow arrow).

**b** Tubules with wider diameters, a tubule attached to a vesicle partially studded with Arf1 (yellow arrow) as well as an apparently capped tubule (pink arrow).

**c** Rectangular Arf1-coated membrane patch (yellow arrow).

**d** Potential starting point of tubulation: Individual Arf1 molecules accumulate on vesicles as a second dense layer on top of the lipid bilayer.

**e-g** Some Arf1-covered tubules adopt a range of diameters, even along one continuous tubule.

**h, i** Cryo-electron micrographs showing tubule branching. The tubules are completely covered by Arf1, also in the fork regions. These images are consistent with the *in cellulo* findings of Stockhammer *et al.*[9] that Arf1 can form a continuous network of tubular structures across the cytoplasm.

**k** Cryo-electron micrographs that show round-shaped ends (yellow arrows) of thoroughly Arf1-coated tubules. Scale bars: 500 Å

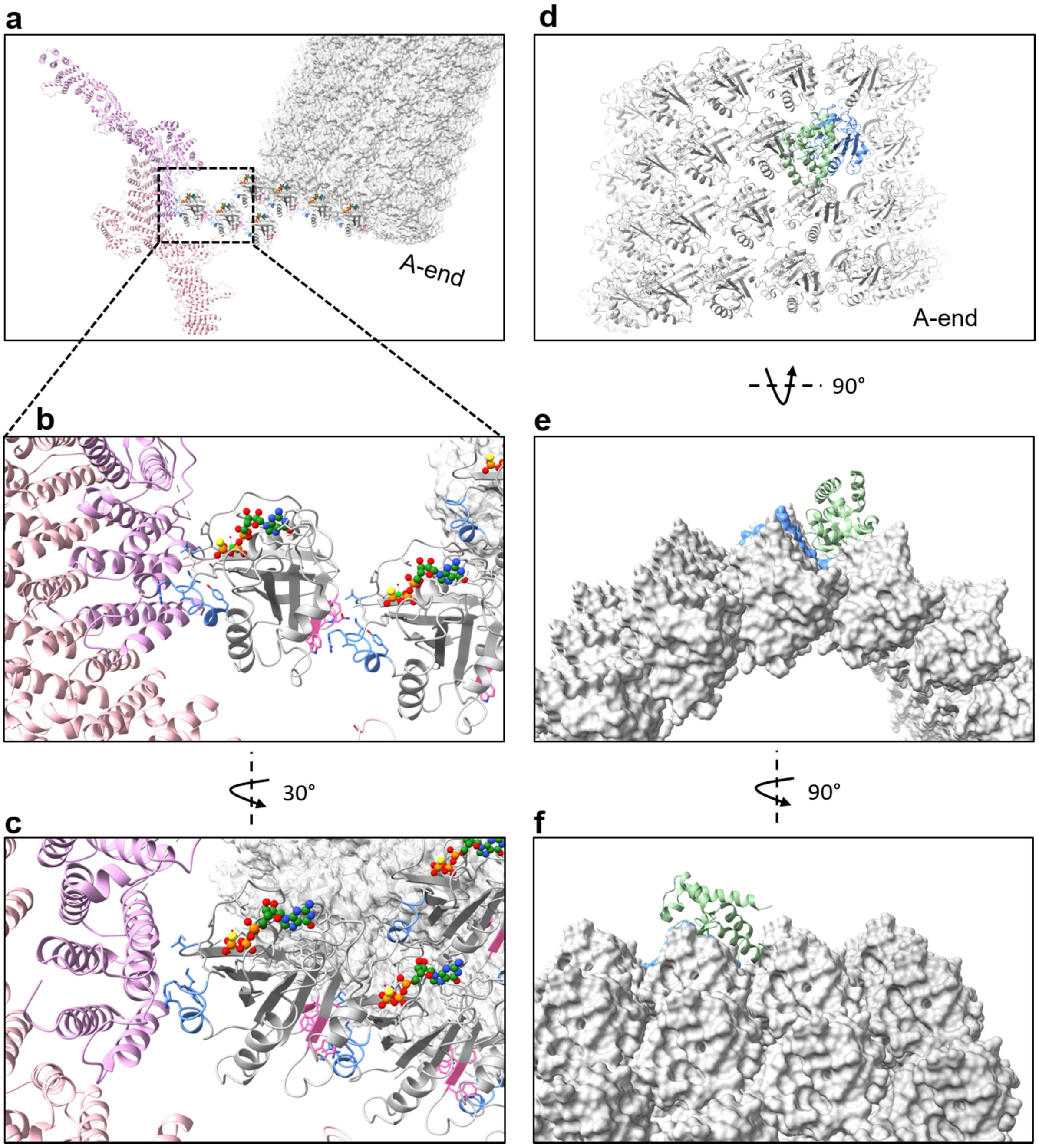


**Supplementary Figure 13:** Interplay of Arf1 with the GEF and GAP effector proteins.

**a-c** Superposition of Arl1 from the Gea2-Arl1 complex (PDB:8ezj)[10] onto Arf1-GTPγS in one filament of the 195 Å tubule (grey cartoon/ surface representation) at the A-end demonstrates that the GEF Gea2 (pale pink) could present activated myr-Arf1-GTP in an orientation suitable for incorporation into the Arf1-coated membrane. Membrane-bound Gea2, which catalyzes the Arf1 GDP-GTP exchange, binds to the switch II region of Arf1 (interface 1', blue) presenting the opposite face (1, pink) in an unobstructed orientation for ready incorporation into the nascent Arf1-GTP tubular helical lattice. The Gea2-catalyzed conversion of soluble Arf1-GDP to membrane-bound Arf1-GTP could therefore have the effect of elongating the membrane-bound Arf1-GTP protofibril.

**b, c** Zoomed-in views of the interaction site between Gea2 and Arf1 switch II region.

**d-f** Superposition of the Arf1 component of the ArfGAP2-Arf1 complex (PDB:5nzs)[11] onto an Arf1-GTPγS G-domain in the Arf1 tubular lattice shows that ArfGAP2 (green ribbons) can dock to a single Arf1 molecule (blue, shown as ribbons (**d**) or in surface representation (**e, f**)) and can access the nucleotide-binding pocket with minimal structural clashes with the tubule.

**e, f** Zoomed-in views of the interaction site.

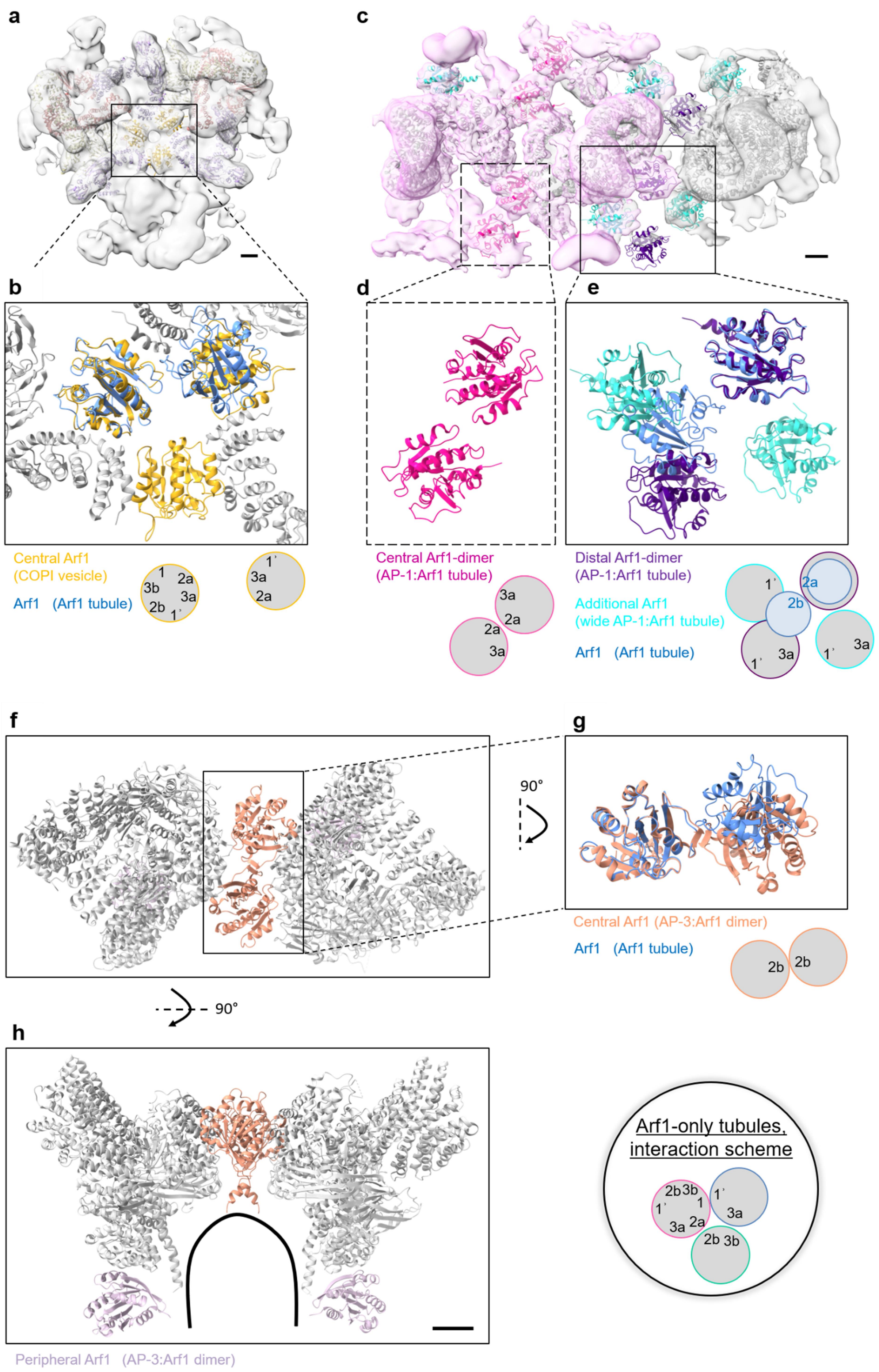
a
b
c
d
e
Central Arf1
(COPI vesicle)
Arf1 (Arf1 tubule)
Central Arf1-dimer
(AP-1:Arf1 tubule)
Distal Arf1-dimer
(AP-1:Arf1 tubule)
Additional Arf1
(wide AP-1:Arf1 tubule)
Arf1 (Arf1 tubule)
f
g
90°
Central Arf1 (AP-3:Arf1 dimer)
Arf1 (Arf1 tubule)
h
Arf1-only tubules, interaction scheme
Peripheral Arf1 (AP-3:Arf1 dimer)

**Supplementary Figure 14**: Comparison of the Arf1-GTPγS tubule arrangement with related Arf1 complex structures.

**a** EM reconstruction of the COPI triad (cryo-EM 3D density map EMD-2985)[12] with corresponding coordinates of the COPI coat proteins (PDB: 5nzr)[11]. To clearly see the triangular arrangement of the membrane-proximal Arf1 molecules, only two asymmetric units (the "leaves") are shown as a ribbon diagram, and the "outer layer" proteins of the coatomer α-COP, ζ-COP, and β-COP have been omitted. Color-coding of the modeled COPI leaf: Arf1 – gold, β'-COP – pale yellow, δ-COP – orange, γ-COP – light purple. Scale bar: 20 Å

**b** Zoomed-in view of (**a**) with the Arf1-GTPγS tubule monomer (blue) superimposed on one of the central Arf1 molecules (gold) in COPI-coated vesicles[11] (individual coatomer proteins shown in grey). The next neighbor in the tubule (along the filament through interface 1) does not align (RMSD of 23.65 Å), nor do any of the other tubule neighbors. In the COPI triad, the Arf1 molecules do not interact with each other directly but rather through interactions with the coatomer proteins.

**c** EM reconstruction of AP-1:Arf1 tubules (cryo-EM 3D density maps EMD-27184 (pale pink) and EMD-27185 (grey)) with corresponding models of the involved AP-1 proteins (PDB: 8d4f and PDB: 8d4g, respectively)[13]. The Arf1 proteins from the AP-1:Arf1 tubule are colored according to Hooy *et al.*[13]: Arf1 in the central dimer (binds to AP-1 β1-subunit), pink; Arf1 in the distal dimer (bridges two AP-1 γ-subunits), dark purple; additional Arf1 molecules (facing towards the γ/σ/μ-subunits of AP-1) appearing in the wider tubes (diameter > 600 Å, compared to EMD-27181/PDB: 8d4c with diameters 200-300 Å from the same study[13]), cyan; adaptor proteins are shown in grey. Scale bar: 20 Å

**d, e** Zoomed-in views of (**c**) show Arf1 dimer interactions within the tubular arrangements found in the AP-1:Arf1 tubules. Overlay with two Arf1 molecules from the Arf1-GTPγS-only tubules (blue) shows the differences in Arf1 G-domain interfaces in both arrangements. Color coding as in (**c**).

**f** The central Arf1 dimer within the dimeric AP-3:Arf1 complex reconstituted in lipid nanodiscs (PDB: 9c59)[14]. As this dimer shows interfaces 2b facing towards each other, it does not align with the Arf1 assembly in the Arf1-coated tubules (RMSD 25.12 Å). AP-3 proteins are shown in grey; Arf1 monomers attached peripherally to AP-3 (in the background) are depicted in pale violet.

**g** Zoomed-in view of the central dimer overlaid with two Arf1 molecules from the Arf1-GTPγS-only tubules (blue).

**h** Side view of the AP-3:Arf1 dimer showing the position of the peripherally bound Arf1 molecules. Color coding as in (**f**). The membrane of the nanodisc is depicted by a solid line. Scale bar: 20 Å. The scheme on the right shows the packing arrangement in Arf1-GTPγS-only tubules (nomenclature according to Arf1 interfaces, compare Fig. 5a).

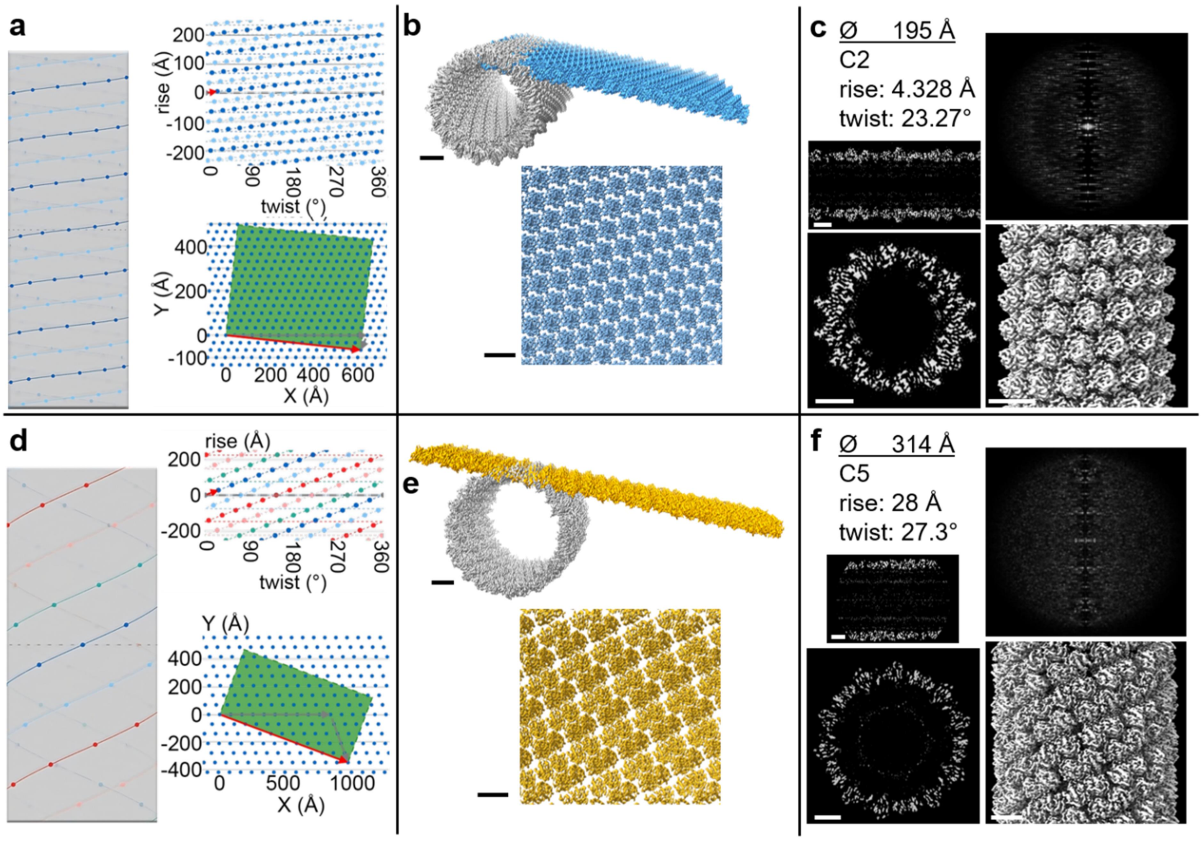


**Supplementary Figure 15**: Comparison of the reconstructed cryo-EM density maps of the Arf1-GTPγS tubules with that of Arf6 tubules (EMD-33414)[15].

**a**, **d** Using the helical parameters of the respective reconstructed cryo-EM density maps, a theoretical 2D crystal lattice was calculated and rolled up to form a 3D helix (web tool https://jiang.bio.purdue.edu/hi3d/)[16]. Distinct differences are apparent between the Arf1 lattice (**a**) and the Arf6 lattice (**d**).

**b**, **e** Unrolling the real-space maps for the Arf1-GTPγS tubules (195 Å, blue (**b**)) and the Arf6 tubules (314 Å, gold (**e**)) demonstrates differences in lattice packing.

**c**, **f** Final helical parameters, power spectra and real space maps of the Arf1-GTPγS (**c**) and the Arf6 tubules (**f**). Cross sections (side and top views) and side views of the tubule are shown. In absence of a power spectrum for the Arf6 tubules, a central slice of the 3D Fourier transformation (FT) of the cryo-EM density map (EMD-33414) (calculated in ChimeraX[17]) is displayed. For comparison, the same procedure was applied to the 195 Å diameter Arf1-GTPγS map. Scale bars: 20 Å

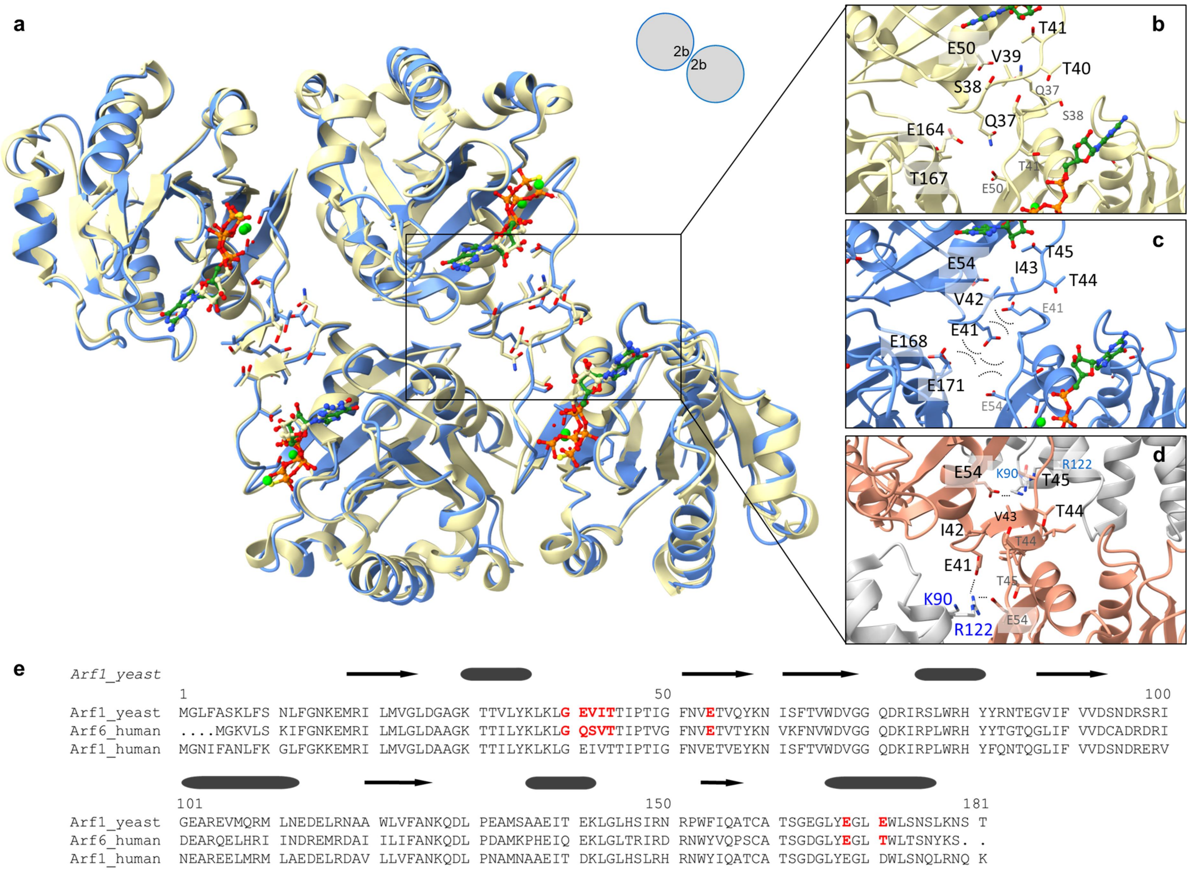


**Supplementary Figure 16**: Superposition of Arf1-GTPγS on the two-fold symmetric Arf6 tetramer of the Arf6 tubules (PDB: 7xrd)[15].

**a** Superposition of four Arf1-GTPγS molecules (blue) upon the Arf6-GTP (light yellow) tetrameric repeat unit (ribbon diagrams with GTPγS or GTP shown in ball-and-stick representation, the magnesium ion shown as a green sphere).

**b** Zoomed-in view of interface Arf6-1 in the Arf6 tubular assembly. Selected residues are labeled (for the second monomer in grey) and shown in stick representation.

**c** Corresponding view of Arf1 in the Arf6 tetrameric arrangement. Replacement of Arf6 residues Q37 and T167 in Arf1 by E41 and E171, respectively, results in the juxtaposition of E41 with its symmetry mate E41' in close proximity to E54 and E168 and E171 of the neighboring Arf1 molecules.

**d** This interface is however observed in the Arf1 dimer in complex with AP-3 (PDB: 9c59, see Supplementary Fig. 15f-h)[14], where E41 and E54' of adjacent Arf1 molecules are juxtaposed by basic residues K90 and R122 of the adaptor protein AP-3.

**e** Sequence alignment of yeast Arf1 with human Arf6 and human Arf1. The secondary structure assignments are derived from the Arf1-GTP$\gamma$S structure. Residues highlighted in red refer to those discussed in panels (**b**) and (**c**).

**Supplementary Movie 1:** Confocal imaging time series of the tubulation reaction of the variant myr-Arf1-R149W$_{2a}$.

**a** A GUV in the center of the image was tubulated by myr-Arf1-R149W$_{2a}$, leading to its explosion into membrane tubules. The prominent fluorescence can be attributed to a bundle of tubules.

**b** Close inspection of further GUVs reveals short, rigid tubules emanating from the GUVs that slowly grow in length (indicated by a red arrow).

A non-linear intensity scale ($\gamma = 0.45$) was used and image contrast was increased to discern the tubular extensions.

**Supplementary Table 1**: Cryo-EM data collection, reconstruction, and helical symmetry parameters.

| | tube diameter 195 Å<br>PDB: 8S5C<br>EMD-19731 | tube diameter 215 Å<br>PDB: 8S5D<br>EMD-19732 |
|---|---|---|
| **Microscopy** | | |
| Magnification | 92,000 × | |
| Voltage (kV) | 200 | |
| Focal length (mm) | 3.4 | |
| Cs (mm) | 2.7 | |
| Objective Aperture (µm) | 70 | |
| Number of movies | 12,483 | |
| total electron dose (e-/Å²) | 50.0 | |
| Defocus range (µm) | -0.6 to -2.6 | |
| Pixel size (Å) | 1.5304 | |
| **Processing** | | |
| No. of symmetries tested | 6 | 32 |
| *Helical symmetry parameters* | | |
| Symmetry imposed | C2 | C3 |
| Rise (Å) | 4.328 | 5.821 |
| Twist (°) | 23.27 | 20.78 |
| Number of subunits per helical turn | 15.47 | 17.32 |
| Pitch (Å) | 66.96 | 100.83 |
| **Particle and map statistics** | | |
| Initial particle images (no.) | 1,489,051 | 1,489,051 |
| Final particle images (no.) | 127,727 | 27,285 |
| Map resolution (Å) | 3.1 | 3.8 |
| FSC threshold | 0.143 | 0.143 |
| Map resolution range (Å) | 3.1 - 6.0 | 3.8 - 7.2 |
| Map sharpening *B*-factor (Å²) | -108.5 | -74.2 |

**Supplementary Table 2**: Statistical analysis of the HADDOCK results of Arf1-Arf1 interaction sites in the tubular arrangement. Cluster size was 20.

| | Interface 1 | Interface 2 | Interface 3 |
|---|---|---|---|
| van der Waals energy (kcal/mol)[a] | -45.1 ± 2.9 | -13.9 ± 1.4 | -19.8 ± 1.1 |
| electrostatic energy (kcal/mol)[a] | -47.9 ± 9.2 | -42.4 ± 8.6 | -169.2 ± 17.2 |
| desolvation energy (kcal/mol)[a] | -13.3 ± 1.7 | 11.7 ± 2.5 | 18.7 ± 3.9 |
| HADDOCK score (a.u.)[a] | -68.0 ± 2.1 | -10.7 ± 2.0 | -34.9 ± 5.1 |
| buried surface area (Å$^2$)[a] | 1028.1 ± 29.5 | 546.4 ± 17.9 | 813.5 ± 26.1 |

[a] Values are given with their standard deviation (1 σ).

**Supplementary Table 3**: Statistical analysis of the HADDOCK results of Arf6-Arf6 interaction sites within the C2 symmetric dimer. Cluster size was 20.

| Arf6- | Interface 1 | Interface 2 | Interface 3 |
|---|---|---|---|
| partly corresponding to Arf1- | int2' | int3' | int1, int2 |
| van der Waals energy (kcal/mol)[a] | -19.9 ± 0.7 | -27.9 ± 3.2 | -57.4 ± 1.9 |
| electrostatic energy (kcal/mol)[a] | -25.1 ± 2.6 | -218.5 ± 15.6 | -67.3 ± 16.8 |
| desolvation energy (kcal/mol)[a] | 26.0 ± 3.6 | 30.8 ± 1.1 | -6.3 ± 1.3 |
| HADDOCK score (a.u.)[a] | 1.1 ± 3.0 | -40.8 ± 4.6 | -77.2 ± 1.0 |
| buried surface area (Å$^2$)[a] | 592.0 ± 6.4 | 1012.8 ± 21.2 | 1547.2 ± 42.9 |

[a] Values are given with their standard deviation (1 σ).

**Supplementary Table 4**: DNA primers used for mutagenesis.

| Name | Sequence (5’ to 3’) |
| --- | --- |
| R149Afw | GGTTTACATTCTATT GCC AACCGTCCATGG |
| R149Arv | CCATGGACGGTT GGC AATAGAATGTAAACC |
| R149Wfw | GGTTTACATTCTATTTGGAACCGTCCATGG |
| R149Wrv | CCATGGACGGTTCCAAATAGAATGTAAACC |
| R149Yfw | GGTTTACATTCTATTTACAACCGTCCATGG |
| R149Yrv | CCATGGACGGTTGTAAATAGAATGTAAACC |
| Arf1pY35Afw | GACCACCGTTTTGGCCAAGTTGAAATTGG |
| Arf1pY35Arv | CCAATTTCAACTTGGCCAAAACGGTGGTC |
| Q57Efw | CGTTGAAACTGTCGAATATAAGAACATTTC |
| Q57Erv | GAAATGTTCTTATATTCGACAGTTTCAACG |
| D114Nfw | GTTGAACGAAAACGAATTGAGAAACGC |
| D114Nrv | GCGTTTCTCAATTCGTTTTCGTTCAAC |
| Q57Nfw | CGTTGAAACTGTCAACTATAAGAACATTTC |
| Q57Nrv | GAAATGTTCTTATAGTTGACAGTTTCAACG |
| D114Efw | GTTGAACGAAGAAGAATTGAGAAACGC |
| D114Erv | GCGTTTCTCAATTCTTCTTCGTTCAAC |

**Supplementary Table 5**: Modelling statistics.

| | tube diameter 195 Å<br>PDB: 8S5C<br>EMD-19731 | |
|---|---|---|
| **Refinement** | | |
| Initial model used (PDB code) | 2ksq | |
| Model resolution (Å) | 3.1 | |
| FSC threshold | 0.5 | |
| Model resolution range (Å) | | |
| Map sharpening *B*-factor (Å²) | 108.5 | |
| *Model composition* [a] | | |
| No. of atoms (protein) | 2609 | |
| No. of atoms (ligands) | 52 | |
| No. of atoms (total) | 2661 | |
| *B-factor (Å²)* | | |
| Protein | 40.88 | |
| Ligand | n.a. | |
| R.m.s. deviations | | |
| Bond lengths (Å) | 0.01 | |
| Bond angles (°) | 0.748 | |
| **Structure validation** | | |
| MolProbity score | 1.56 | |
| Clash score | 2.53 | |
| Rotamer correctness (%) | 96.4 | |
| *Ramachandran plot* | | |
| Favoured (%) | 97.52 | |
| Allowed (%) | 2.48 | |
| Disallowed (%) | 0 | |
| **Data** | | |
| *Resolution estimates (Å)* | Masked | Unmasked |
| d FSC (half maps, 0.143) | --- | --- |
| d 99 (full/half1/half2) | 3.3/---/--- | 3.2/---/--- |
| d model | 3.2 | 3.3 |
| d FSC model (0/0.143/0.5) | 2.9/3.0/3.1 | 3.0/3.1/3.2 |

| | |
|---|---|
| Map min/max/mean | -52.00/94.82/0.66 |
| Model vs. Data | |
| CC (mask) | 0.90 |
| CC (box) | 0.65 |
| CC (peaks) | 0.62 |
| CC (volume) | 0.86 |
| Mean CC for ligands | 0.85 |

[a] Regarding one protein subunit, i.e., one Arf1 molecule and its ligands.
